# A novel wild-derived MHC-linked locus regulates host immunity to gammaherpesvirus infection

**DOI:** 10.64898/2026.06.23.733957

**Authors:** Courtney M. Waytashek, Emily A. Nelson, Katherine J. Sessions, Kaylin Dalhberg, Meghan Fondakowski, Jason A. Bubier, Edward J. Usherwood, Dimitry N. Krementsov

## Abstract

Inefficient host control of infection by gammaherpesviruses is a risk factor for lymphoproliferative disease, autoimmunity, and cancer, yet naturally occurring genetic determinants of viral control remain poorly defined. Previously, we discovered that wild-derived PWD/PhJ (PWD) mice exhibit markedly improved control of murine gammaherpesvirus 68 (MHV-68) replication compared with C57BL/6J (B6) mice. This elite control of viral replication was linked to muted T cell responses, but partially dependent on NK cells. Here, we used genetic approaches to identify host determinants controlling MHV-68 resistance in PWD mice. Analysis of B6PWDF1 and B6PWDF2 populations suggested the existence of a major locus in the PWD genome contributing to MHV-68 resistance. Using B6.Chr^PWD^ chromosome substitution (consomic) mice, we mapped a major resistance locus to Chr 17 and refined the required interval to 27.6-49.4 Mb, including the MHC locus. While B6.Chr17^PWD^ mice recapitulated muted T cell responses observed in PWD mice, they controlled viral burden independent of NK cells and without reduced frequencies of infected germinal center B cells, indicating that resistance of PWD mice comprises multiple genetically and mechanistically distinct pathways. While the NK cell receptor complex on PWD chromosome 6 did not provide protection, genotype-phenotype analysis of the B6PWDF2 cohort revealed additional non-Chr 17-linked loci contributing to viral control. Together, our results identify a novel MHC-linked locus regulating gammaherpesvirus burden with minimal cytotoxic T cell expansion, demonstrating that effective host control of gammaherpesviruses can be achieved by diverse immune mechanisms.

**Importance:** Poor host control of human gammaherpesvirus infection is linked to cancer and autoimmunity, but how natural genetic variation impacts immune mechanism of control is poorly defined. In this study, we examined genetic mechanisms linked to superior control of gammaherpesvirus infection. We identified a novel MHC-linked locus that regulates gammaherpesvirus burden without a large expansion of cytotoxic T cells and independent of NK cells. These findings show that host control of gammaherpesviruses can be achieved through diverse immune mechanisms that are driven by variation in host genetic background.

## Introduction

Epstein-Barr virus (EBV) and Kaposi’s sarcoma-associated herpesvirus (KSHV) are lymphotropic herpesviruses in the gammaherpesvirus subfamily(1, 2). Both viruses are oncogenic(2, 3). Additionally, EBV infection is associated with an increased risk of autoimmune disease, including multiple sclerosis (MS)(4, 5), rheumatoid arthritis(6), and systemic lupus erythematosus(7–9). EBV is an enveloped virus with a double-stranded DNA genome that has lytic and latent replication as two distinct phases of the viral lifecycle, which results in persistent life-long infection of the host(1, 2). EBV is primarily transmitted via saliva, although it can also be transferred via organ transplantation, blood transfusion, or through sexual contact(10). EBV establishes lytic infection in epithelial cells and latent infection in memory B cells, which disseminate the virus to other lymphoid organs throughout the body. EBV infects over 90% of the global population, with most infections occurring asymptomatically in the first 1-2 years of life(1, 6). When infection is delayed until adolescence, the risk of symptomatic infection in the form of infectious mononucleosis (IM) increases(1). IM is an immune-driven syndrome which is characterized by persistent fatigue, fevers, and splenomegaly that can last for weeks to months.

The outcomes of EBV infection in individuals with primary immunodeficiencies have highlighted the need for immunological control of viral replication during the acute phase, otherwise life-threatening conditions such as chronic active EBV, hemophagocytic lymphohistiocytosis, and uncontrolled IM occur(2, 11, 12). Similarly, secondary immunodeficiencies can result in EBV reactivation and EBV-associated malignancies such as post-transplant lymphoproliferative disease(13). Furthermore, there is evidence in healthy individuals that poor control of EBV, either as a diagnosis of IM, high antibody titers against EBV antigens, or high EBV viral load, is associated with elevated risk of developing MS and some EBV-associated malignancies, such as Hodgkin’s lymphoma(13–15). Across these conditions, inadequate host control of EBV replication emerges as a unifying driver of pathology.

Initial genome-wide association studies evaluating indirect measures of the severity of EBV infection have suggested that poor viral control is genotype dependent(14, 15). However, the underlying mechanisms behind how genetic variation impacts the immunological response to EBV and efficiency of host control of viral replication, are not known. Furthermore, mechanistic study of primary infection with EBV in humans is particularly challenging, due to the persistent and often asymptomatic nature of infection. To overcome this, human gammaherpesvirus infection can be modeled in mice using murine gammaherpesvirus-68 (MHV-68), a murine gammaherpesvirus related to KSHV and EBV, allowing for in-depth mechanistic studies(16).

However, a limitation of the MHV-68 model is that it has been primarily applied to conventional laboratory inbred mice like C57BL/6 (B6) and BALB/c, which, unlike humans, capture limited genetic and phenotypic diversity. Our lab has recently shown that infection of the genetically divergent wild-derived mouse strain PWD/PhJ (PWD) results in elite viral control of MHV-68 replication, with viral loads several logs lower than B6 mice during acute infection and latency(17). We further showed that B6 mice exhibit expansion of a unique CX3CR1+ KLRG1+ CD4+ cytotoxic helper T cell (ThCTL) subset during chronic MHV-68 infection, while this subset remained absent in the resistant PWD mice(17). In contrast, PWD mice exhibited a novel NK cell dependency for efficient control of MHV-68 replication, with an increase in splenic viral burden seen after NK cell depletion in PWD, but not B6 mice(17), consistent with the essential role for NK cells in control of EBV in humans(2, 11, 12).

The aim of this study was to investigate host genetic factors contributing to MHV-68 resistance in PWD mice and determine the underlying immunological mechanisms. We found evidence of multiple loci in the PWD genome contributing to control of MHV-68 replication, with a major dominant locus identified on chromosome 17 between 27.6-49.4 Mb. Notably, this locus contains the entirety of the MHC/*H2* region. A consomic strain carrying the full PWD chromosome 17, Chr17^PWD^, captured partial resistance to MHV-68, recapitulated a PWD-like lack of expansion of ThCTL and effector CD8 T cells, but did not require NK cells for control of viral burden, suggesting multiple independent genetic mechanisms of resistance in PWD mice. While the NK cell receptor locus (NKC) on Chr6 was not sufficient for resistance, genetic analysis of B6PWDF2 mice suggested the existence of additional loci. Together, our findings identify a novel MHC-linked locus on PWD chromosome 17 that regulates control of gammaherpesvirus viral burden without an expansion of cytotoxic T cells or the requirement for NK cells, suggesting that gammaherpesvirus is controlled by diverse host genetic mechanisms.

## Results

### Intercrosses between B6 and PWD mice demonstrate the existence of a major locus driving MHV-68 resistance

In our previous study (17), we reported that the wild-derived strain PWD/PhJ (PWD) is an elite controller of MHV-68 replication in lymphoid tissue, compared to B6 mice. This suggested the existence of genetic variant(s) in the PWD genome that enhance host control of MHV-68 replication. To assess the inheritance pattern of the resistance phenotype, we crossed B6 and PWD mice to generate a heterozygous B6PWDF1 population. B6PWDF1 mice, together with B6 and PWD parental controls, were infected with MHV-68 via the intraperitoneal (i.p.) route and sacrificed on day 9 post infection, the timepoint for peak viral replication in the spleen, when the largest difference in viral load between B6 and PWD mice was previously observed (17). We found that B6PWDF1 mice had a significant (∼5 fold) reduction in MHV-68 viral DNA load in the spleen compared to B6 mice, an intermediate reduction compared with PWD mice (**Fig. 1A**). These results suggested several plausible inheritance patterns for our trait, including incomplete dominance at a major locus, or additive inheritance of multiple loci in the PWD genome that enhance host control of MHV-68.

**Figure 1.**
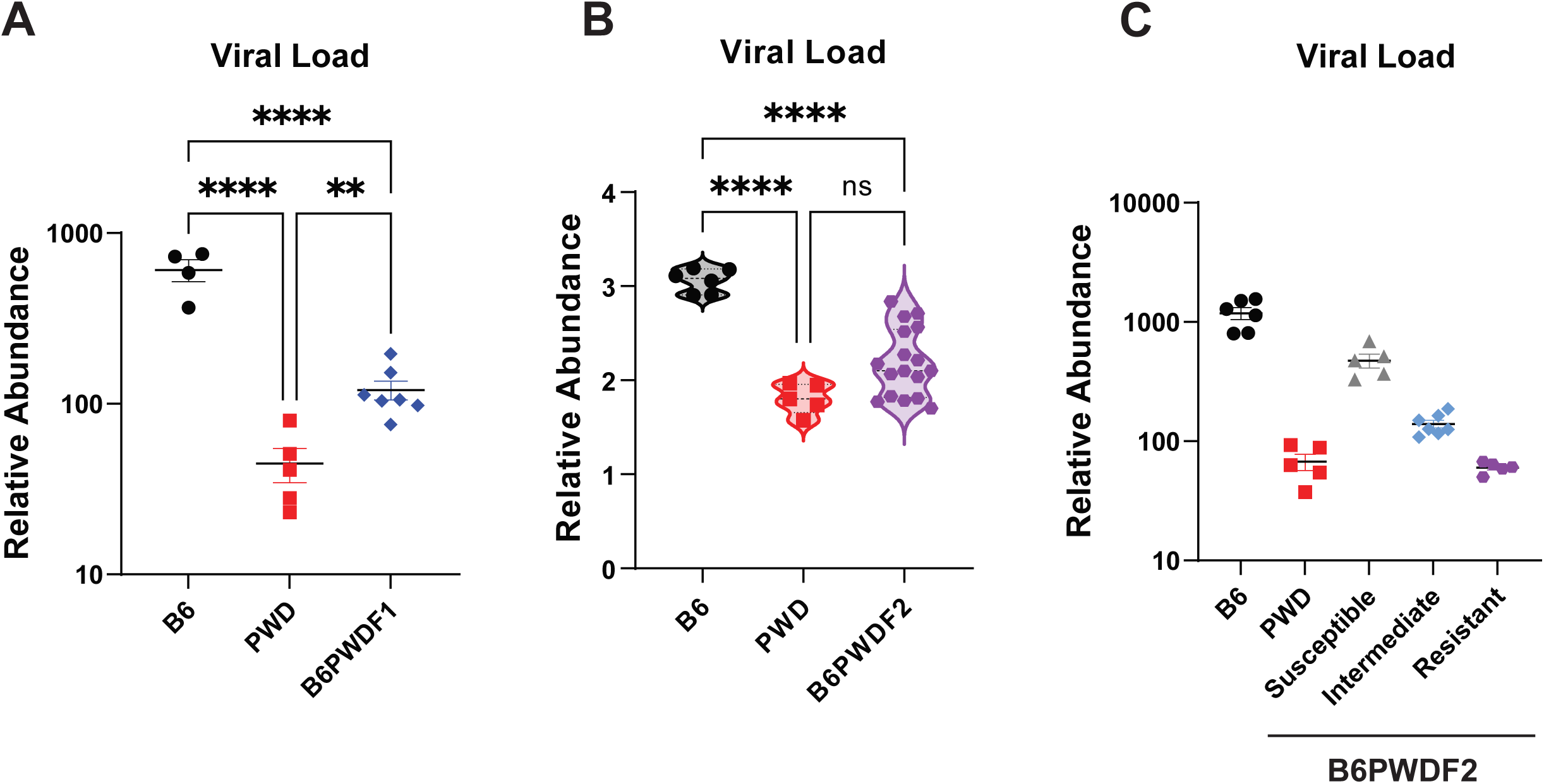
B6 × PWD intercrosses suggest the existence of a single major locus of MHV-68 resistance in the PWD genome. A cross between B6 (F) and PWD (M) mice was completed to generate a B6PWDF1 population. B6PWDF1 mice were then crossed to generate a B6PWDF2 population. Male and female F1 (**A**) and F2 (**B-C**) offspring from the cross and B6 and PWD controls were infected with 105 PFU of MHV-68 via the i.p. route. At d9 post-infection, spleens were collected and replication of MHV-68 was assessed by measuring relative abundance of MHV-68 ORF50 DNA by qPCR. Data in (**C**) represents the same data presented in (**B**) with B6PWDF2 mice stratified into susceptible, intermediate, and resistant groups based on DNA viral load. Grouping was done as follows: susceptible: >200, intermediate: 200-100, and resistant: <100. Significance of differences between genotypes was determined in (**A**) and (**B**) by lognormal one-way ANOVA with Tukey’s multiple comparisons test. For (**A**) and (**B**): ** indicates p≤0.01, *** indicates p≤0.001, and **** indicates p≤0.0001.

To further test the mode of inheritance, we performed an intercross of B6PWDF1 mice to generate a small B6PWDF2 population with randomly occurring recombination events throughout their genomes. B6PWDF2 mice, along with B6 and PWD parental controls, were infected i.p. with MHV-68 and sacrificed on day 9 post infection. When we assessed splenic viral load, we found that the B6PWDF2 mice displayed a trimodal distribution, with one group demonstrating complete resistance, another group F1-like intermediate resistance, and a third group that was B6-like (**Fig. 1B and C**). This trimodal distribution is most consistent with the presence of a single major locus of resistance that shows incomplete dominance, although it does not rule out the presence of a major locus with modifiers.

### Genetic mapping of MHV-68 resistance identifies a major locus on PWD chromosome 17

To map these variants to a particular PWD chromosome, we utilized a partial panel of B6.Chr^PWD^ chromosome substitution (consomic) mice (18). These consomic strains contain a single homozygous PWD chromosome pair on a B6 inbred strain background. At the time of the experimental work, a subset of 8 strains of the complete 28-strain B6.Chr^PWD^ consomic panel were available in our vivarium and were included in the screen. Consomic mice, together with B6 and PWD parental controls, were infected i.p. with MHV-68 and sacrificed on day 9 post infection. Spleens were collected and viral replication was assessed by measuring relative abundance of viral DNA via qPCR (viral load), or infectious viral titer by plaque assay. The majority (7/8) of consomic strains tested were not significantly different from B6 by either measure of viral replication. However, the strain B6.Chr17^PWD^ (Chr17^PWD^) displayed a 4-fold reduction in viral load compared to B6 mice, a partial reduction compared with the 18-fold reduction in viral titer in PWD mice (**Fig. 2A**). Strikingly, the viral titer in Chr17^PWD^ mice was reduced to undetectable levels, thus fully recapitulating the PWD resistance phenotype by this measure (**Fig. 2B**). These results demonstrate that a major MHV-68 resistance locus present in the PWD genome maps to chromosome 17 (Chr 17).

**Figure 2.**
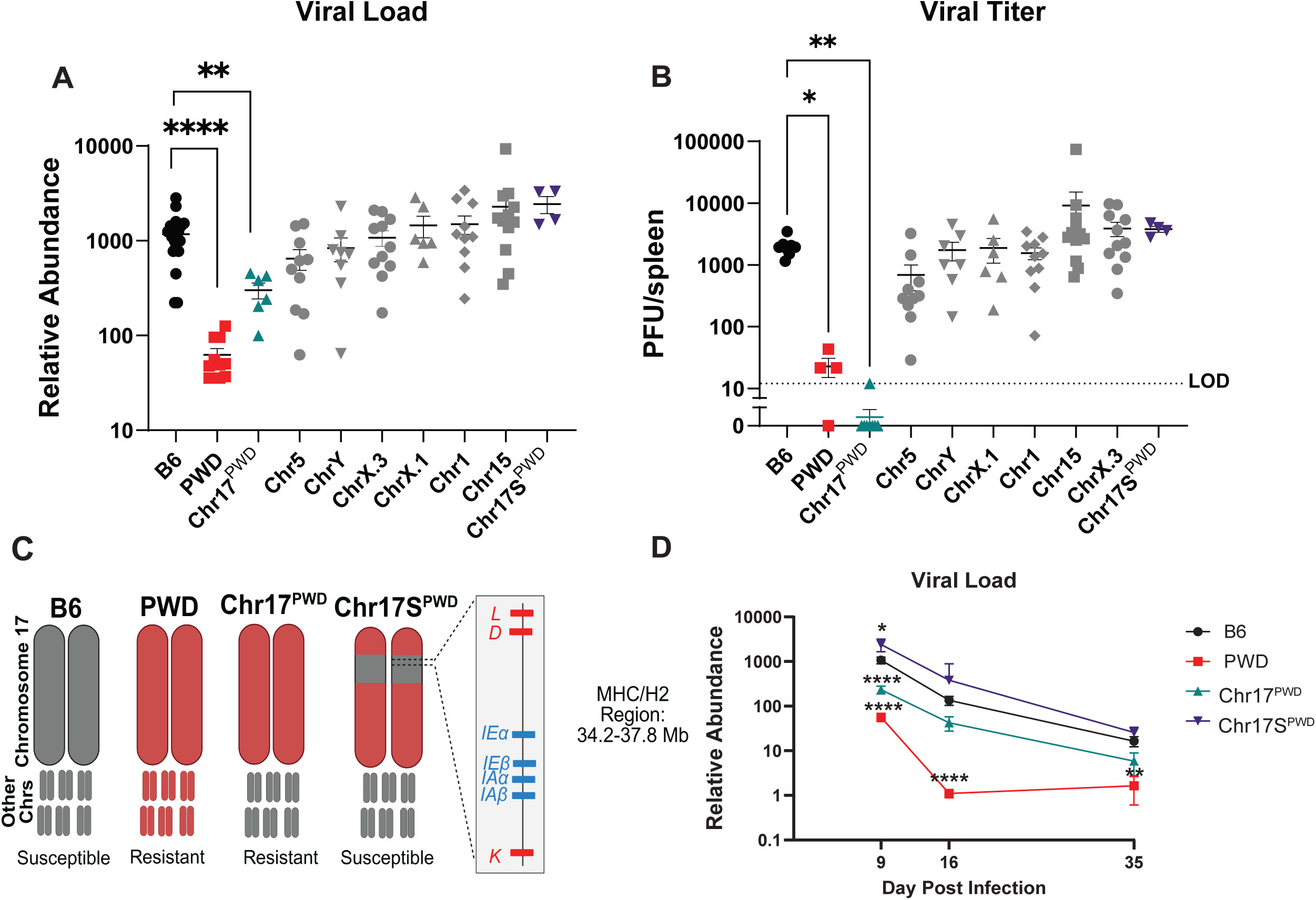
Mapping MHV-68 resistance using B6.ChrPWD consomic mice. (**A and B**) Male and female B6.ChrPWD consomic mice and B6 and PWD controls were infected with 105 PFU of MHV-68 via the i.p. route. At days 9, 16, and 35 post-infection, spleens were collected and replication of MHV-68 was assessed. **(A**) Relative abundance of MHV-68 DNA load, as assessed by qPCR for the ORF50 gene at day 9. (**B**) Infectious viral titers (expressed as plaque forming units (PFU) per spleen) at day 9. Data in (**A**) and (**B**) represent both sexes, pooled from two and three independent cohorts of mice respectively. Significance of differences between genotypes in (**A**) were determined by one-way ANOVA with Dunnett’s multiple comparisons test on the log transformed data. Significance of differences between genotypes in (**B**) were determined by a Kruskal-Wallis test. (**C**) Diagram depicting chromosome 17 and other chromosomes in B6, PWD, Chr17PWD, and Chr17SPWD mice, highlighting the homozygous B6-derived region on chromosome 17 for the Chr17SPWD strain and the location of the MHC/H2 region. (**D**) Relative abundance of MHV-68 DNA load, as assessed by qPCR for the ORF50 gene at days 9, 16, and 35. Each timepoint (9, 16, 35) are from independent cohorts of mice, with three experiments pooled for d9 data, shown with sexes pooled. Cohort sizes for each timepoint were as follows; D9: B6=14, PWD=8, Chr17PWD=9, Chr17SPWD=9; D16: B6=5, PWD=6, Chr17PWD=4, Chr17SPWD=6; D35: B6=9, PWD=4, Chr17PWD=6, Chr17SPWD=6. Significance of differences between genotypes in (**D**) were determined by one-way ANOVA on the log-normalized data for each independent time point with Dunnett’s multiple comparisons test. For (**A**), (**B**), and (**D**): * indicates p≤0.05, ** indicates p≤0.01, *** indicates p≤0.001, and **** indicates p≤0.0001.

Notably, our consomic panel included two strains covering PWD Chr 17. While the Chr17^PWD^ consomic strain carries the entire PWD Chr 17, the B6.Chr17S^PWD^ sub-consomic strain (designated Chr17S^PWD^; “S” for short) carries a homozygous B6-derived interval in the middle of Chr 17, with the two breakpoints that we determined in our previous studies to be located between 26 and 34.3 Mb and 45.5 and 47.7 Mb, respectively (19). Here, we used high resolution genotyping (see materials and methods) to more precisely define these breakpoints. We found that the B6-derived locus in Chr17S^PWD^ mice encompassed a region located between 27.6 – 49.4 Mb on Chr 17, which includes the entire MHC/*H2* locus at ∼34.2-37.8 Mb (**Fig. 2C**). Importantly, unlike the full-length Chr17^PWD^ mice, Chr17S^PWD^ sub-consomic mice completely lacked resistance to MHV-68 (**Fig. 2A** and **2B**). Given the genetic difference between these two strains, these results demonstrate the MHV-68 resistance phenotype controlled by PWD chromosome 17 requires the variants encoded in the region between 27.6-49.4 Mb.

We next asked if Chr17^PWD^ and Chr17S^PWD^ mice retain their respective resistant and susceptible phenotypes over the course of MHV-68 infection. B6, PWD, Chr17^PWD^, and Chr17S^PWD^ mice were infected i.p. with MHV-68 and spleens were collected at days 9, 16, and 35 post infection, to assess peak lymphoid tissue replication, early latency, and full latency, respectively. As above, at d9 post infection, Chr17^PWD^ mice had significantly reduced (4.5-fold) splenic viral loads compared to B6 mice, while Chr17S^PWD^ mice in fact exhibited significantly higher viral loads than B6 controls (**Fig. 2D**). The same pattern (2–3-fold reduction in Chr17^PWD^ relative to B6) was observed for d16 and d35 post-infection, albeit the difference between consomic strains and B6 controls was not significant, and Chr17^PWD^ mice did not mirror the sharp decrease in detectable viral DNA observed in PWD mice at d16 (**Fig. 2D**). Taken together, these results indicate that the trends in strain-specific differences in early control of viral replication are maintained throughout the course of infection.

The natural route of infection for MHV-68 remains debated, but in addition to i.p., a common route of experimental infection is via the intranasal route. To ensure our findings were not dependent on the route of infection, we also performed intranasal infections of B6, PWD, and Chr17^PWD^ mice, assessing viral replication at days 7 and 35 post-infection. During acute replication in the lungs, we found no difference in viral load between the strains (**Supplemental Figure S1A**). However, in the spleen, compared to B6 controls, PWD mice had significantly reduced viral load at both timepoints, while Chr17^PWD^ mice had significantly lower viral load at the early d7 timepoint and 8-fold non-significant reduction at d35 (**Supplemental S1B** and **S1C**). These kinetics in the spleen mirror those observed above with i.p. infection, suggesting that PWD and Chr17^PWD^ mice exhibit enhanced control of early lymphoid tissue viral replication, independent of the route of infection.

### Chr 17-driven MHV-68 resistance is associated with muted cytotoxic T cell responses

Next, we used flow cytometry to examine the kinetics of splenic immune responses in our four strains of interest at d9, d16, and d35 post-MHV-68-infection (i.p.). We first assessed the frequency of major lymphocyte populations between B6 controls and strains. We found numerically minor but statistically significant differences between B6 and PWD and/or the two consomic strains in the frequency of B cells, CD4+ T cells, CD8+ T cells, and NK cells, at individual time points (**Fig. 3A-D**). However, these differences did not segregate with the MHV-68 resistance profiles of these strains, e.g., some changes were unique to PWD, other changes were shared between Chr17^PWD^ and Chr17S^PWD^, suggesting that enhanced host control of viral burden driven by Chr 17 is not associated with changes in major lymphocyte subsets.

**Figure 3.**
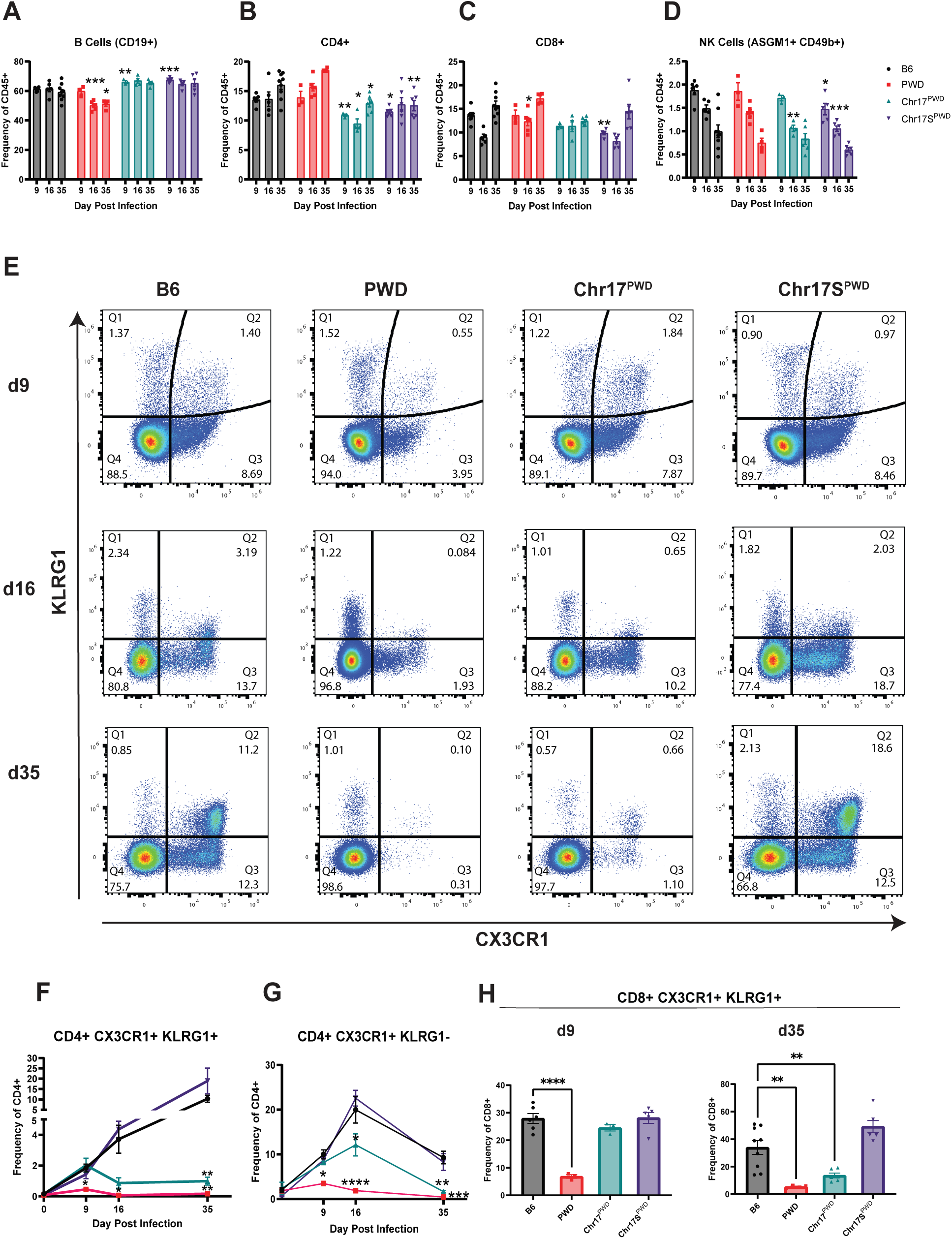
MHV-68-resistant Chr17PWD and PWD mice lack an expansion of cytotoxic effector T cells in the spleen during MHV-68 infection. Male and female B6, PWD, Chr17PWD, and Chr17SPWD mice were infected with 105 pfu of MHV-68 via the i.p. route. Spleens were collected at days 9, 16, and 35 and flow cytometry was completed to assess immune populations. Each timepoint (9, 16, 35) are from independent cohorts of mice, with three experiments pooled for d9 data, all shown with sexes pooled. Cohort sizes for each timepoint were as follows; D9: B6=14, PWD=8, Chr17PWD=9, Chr17SPWD=9; D16: B6=5, PWD=6, Chr17PWD=4, Chr17SPWD=6; D35: B6=9, PWD=4, Chr17PWD=6, Chr17SPWD=6. (**A-D**) Frequency of B cells (CD19+), CD4+ T cells (TCRβ+ CD4+), CD8+ T cells (TCRβ+ CD4+), and NK cells (ASGM1+ CD49b+), calculated as percent of CD45+ leukocytes. (**E**) Representative flow cytometric plots of CX3CR1 and KLRG1 expression by CD4+ T cells on days 9, 16, and 35 post infection. (**F-H**) Frequency of CX3CR1+ KLRG1+ CD4+ T cells, CX3CR1+ KRLG1- CD4+ T cells (calculated as percent of CD4+ T cells), and CX3CR1+ KLRG1+ CD8+ T cells (calculated as percent of CD8+ T cells), respectively. Significance of differences between B6 controls and the other 3 strains in (**A-D, F-G**) were determined for each timepoint (9, 16, 35) independently by one-way ANOVA with Dunnett’s multiple comparisons test, except for d16 and d35 in **F** and d35 in **G** (Brown-Forsythe) and d9 in **G** (Kruskal). Significance of differences between B6 controls and the other 3 strains in (**H**) were determined independently for d9 by one-way ANOVA with Dunnett’s multiple comparisons test and d35 by Brown-Forsythe and Welch one-way ANOVA with Dunnett’s T3 multiple comparisons test. For (**A-H**): * indicates p≤0.05, ** indicates p≤0.01, *** indicates p≤0.001, and **** indicates p≤0.0001.

In our previous work (17), we found that over the course of MHV-68 infection, B6 mice show a continuous expansion of a unique CX3CR1+ KLRG1+ cytotoxic CD4+ T helper (ThCTL) subset, which is surprisingly absent in PWD mice. We therefore asked if this phenotype segregates with the PWD resistance locus on chromosome 17. We found that while susceptible B6 and Chr17S^PWD^ mice exhibited a continued expansion of CX3CR1+ KLRG1+ CD4+ T cells over the course of MHV-68 infection, this expansion was absent in resistant Chr17^PWD^ and PWD mice (**Fig. 3E** and **F**). We previously demonstrated that the expansion of CX3CR1+ KLRG1+ CD4+ T cells is temporally preceded by an expansion of CX3CR1+ KLRG1- CD4+ T cells, which begins on d9, and peaks at d16 post-infection (17). This was the pattern observed in susceptible B6 and Chr17S^PWD^ mice, while PWD mice largely lacked this population (**Fig. 3E** and **G**). In contrast, Chr17^PWD^ mice exhibited a B6-like expansion of CX3CR1+ KLRG1- CD4+ T cells at d9, which began to contract at d16 and were completely absent by d35 (**Fig. 3E** and **G**). Together, these results suggest that: 1) resistance to MHV-68 by the PWD-derived locus on chromosome 17 occurs mostly in the absence of expansion of ThCTL cells, and 2) the lack of ThCTL expansion requires a PWD-derived region between 27.6-49.4 Mb on chromosome 17.

In our previous study, in MHV-68-infected B6 mice we found an analogous expansion of long-lived effector-like CD8+ T cells (LLEC; (20)) co-expressing CX3CR1 and KLRG1, which was also muted in PWD mice (17). We found that, like the initial expansion of CX3CR1+ KLRG1-CD4+ T cells, these CX3CR1+ KLRG1+ CD8+ T cells initially exhibited a B6-like expansion in Chr17^PWD^ mice at d9, followed by contraction to near PWD levels by d35 (**Fig. 3H**). Taken together, these results demonstrate that gammaherpesvirus resistance driven by the PWD loci/locus on chromosome 17 is paralleled by reduced and transient cytotoxic effector T cell expansion, suggesting that the latter is a consequence of inefficient host control of viral replication, rather than an indicator of robust host immunity, consistent with findings from EBV (21) and MHV-68 (22) infectious mononucleosis.

### PWD, but not Chr17^PWD^, mice exhibit a reduction in infected germinal center B cells

After its spread to secondary lymphoid organs during primary infection, MHV-68 replicates predominantly in B cells, which subsequently act as the primary viral reservoir during latency. We thus asked if the reduced splenic viral load and titer in PWD and Chr17^PWD^ mice results in reduced infection of a particular subset of B cells. B6, PWD, and Chr17^PWD^ mice were infected with a histone 2B-YFP reporter-expressing clone of MHV-68 (23) via the i.p. route and flow cytometry was used to assess the frequencies of infected YFP-expressing splenocytes and key MHV-68 target cell types, at the peak of lymphoid infection at d9. We found that PWD mice had a significantly lower frequency of B cells (CD19+) compared with both B6 and Chr17^PWD^ mice, and significantly fewer of these CD19+ B cells were YFP+ (**Fig. 4A**). Germinal center B cells are a key target of MHV-68 replication during the acute phase of infection, as this virus usurps the ongoing germinal center reaction to increase the number of potential target cells to infect. It is thus possible that reduced viral loads in PWD and or Chr17^PWD^ mice result from impaired germinal center responses. However, we found that PWD mice have a significantly higher frequency of germinal center (GL7+ CD95+) B cells than either B6 or Chr17^PWD^ mice, with the latter two strains exhibiting comparable frequencies (**Fig. 4B**). In contrast, compared with B6, the frequency of infected YFP+ germinal center B cells was significantly lower in PWD mice, but not in Chr17^PWD^ (**Fig. 4B**). These findings suggest that PWD mice mount a robust germinal center response, but the frequency of germinal center B cell infection is lower compared with B6; a phenotype that is not recapitulated in Chr17^PWD^ mice.

**Figure 4:**
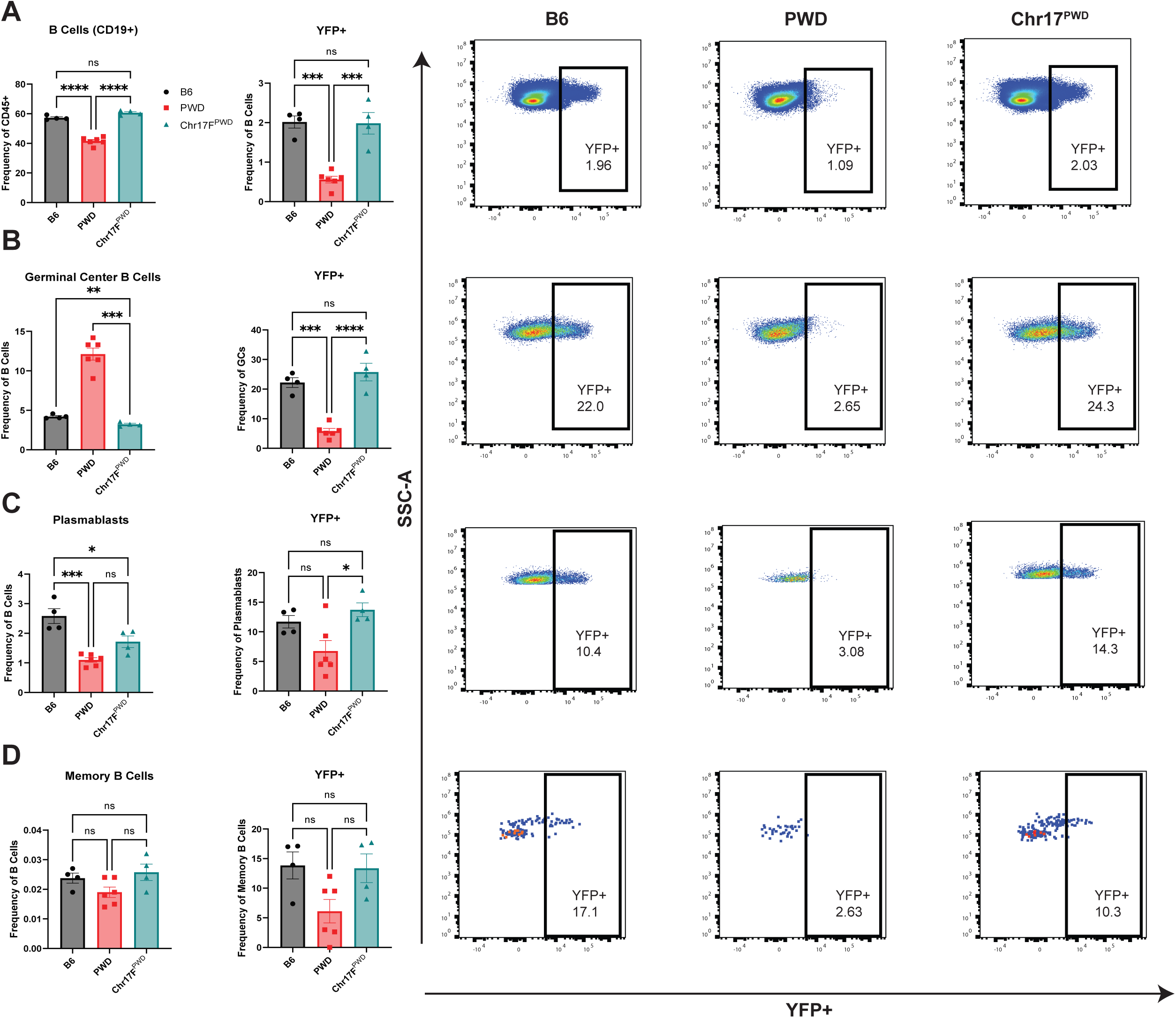
PWD mice have fewer infected B cells. Male and female B6, PWD, and Chr17PWD mice were infected with a YFP reporter-expressing MHV-68 virus via the i.p. route. Spleens were collected at d9 and flow cytometry was performed to assess frequencies of B cell populations and infected YFP+ cells. Male and female data were pooled by genotype. (**A**) Frequency of B cell (CD19+) among CD45+ cells and frequency of YFP+ cells among CD19+ cells. Representative flow of YFP+ B cells. (**B**) Frequency of germinal center B cells (GL7+ CD95+) among CD19+ cells and the frequency of YFP+ cells among germinal center cells. Representative flow of YFP+ germinal center B cells. (**C**) Frequency of plasmablast B cell (B220lo CD138+ SSChigh) among CD19+ cells and the frequency of YFP+ cells among plasmablasts. Representative flow of YFP+ plasmablasts. (**D**) Frequency of memory B cells (B220lo CD80+) among CD19+ cells and the frequency of YFP+ memory B cells. Representative flow of YFP+ memory B cells. Significance of differences between genotypes were determined by one-way ANOVA with Tukey’s multiple comparisons test. Significance of differences between genotypes for (**B, germinal center frequency among CD19+ cells**) were determined by Brown-Forsythe and Welch one-way ANOVA with Dunnett’s T3 multiple comparisons test. For (**A-D**): * indicates p≤0.05, ** indicates p≤0.01, *** indicates p≤0.001, and **** indicates p≤0.0001.

Other key B cell subsets for MHV-68 infection are plasmablasts and memory B cells, the latter a key target for latency. We found that PWD and Chr17^PWD^ mice had significantly fewer plasmablasts (B220^lo^ CD138+ SSC^hi^) compared to B6 mice, and Chr17^PWD^ mice had a significantly higher frequency of YFP+ infected plasmablasts compared with PWD (**Fig. 4C**). Moreover, there was no significant difference between strains in the frequency of memory B cells (B220^lo^ CD80+) or in the frequency of YFP+ infected memory B cells, although the frequency of this population is notably very low at this time point (**Fig. 4D**). Collectively, these results demonstrate that the large reduction in splenic viral load and titer in resistant PWD mice is paralleled by a reduction in the frequency of infected germinal center B cells, a phenotype that is surprisingly absent in resistant Chr17^PWD^ mice.

### Control of viral burden by PWD chromosome 17 is inherited in a completely dominant manner

The partial reduction of viral load in Chr17^PWD^ mice (**Fig. 2A**) contrasted our findings above in B6PWDF2 mice, which had initially suggested the existence of a single partially dominant locus - because if the major locus on Chr 17 represented the single incompletely dominant resistance locus, we would have expected the consomic (homozygous) Chr17^PWD^ mice to fully recapitulate the resistance seen in PWD mice. Thus, in order to determine the pattern of inheritance for our identified locus on chromosome 17, we next crossed B6 and Chr17^PWD^ mice (using reciprocal crosses) to generate heterozygous F1 offspring, which were tested for susceptibility to MHV-68 infection. Regardless of the direction of the cross, B6Chr17^PWD^F1 and Chr17^PWD^B6F1 mice had significantly reduced viral load at d9 post-infection compared with B6 mice, fully recapitulating the partial reduction in viral load seen in homozygous Chr17^PWD^ compared with PWD mice (**Fig. 5A**). Similarly, the B6Chr17^PWD^F1 mice fully recapitulated the reduction in viral titer in Chr17^PWD^, down to a level comparable with PWD (**Fig. 5B**). These results indicate that the enhanced control of MHV-68 burden by loci on chromosome 17 is a completely dominant trait.

**Figure 5.**
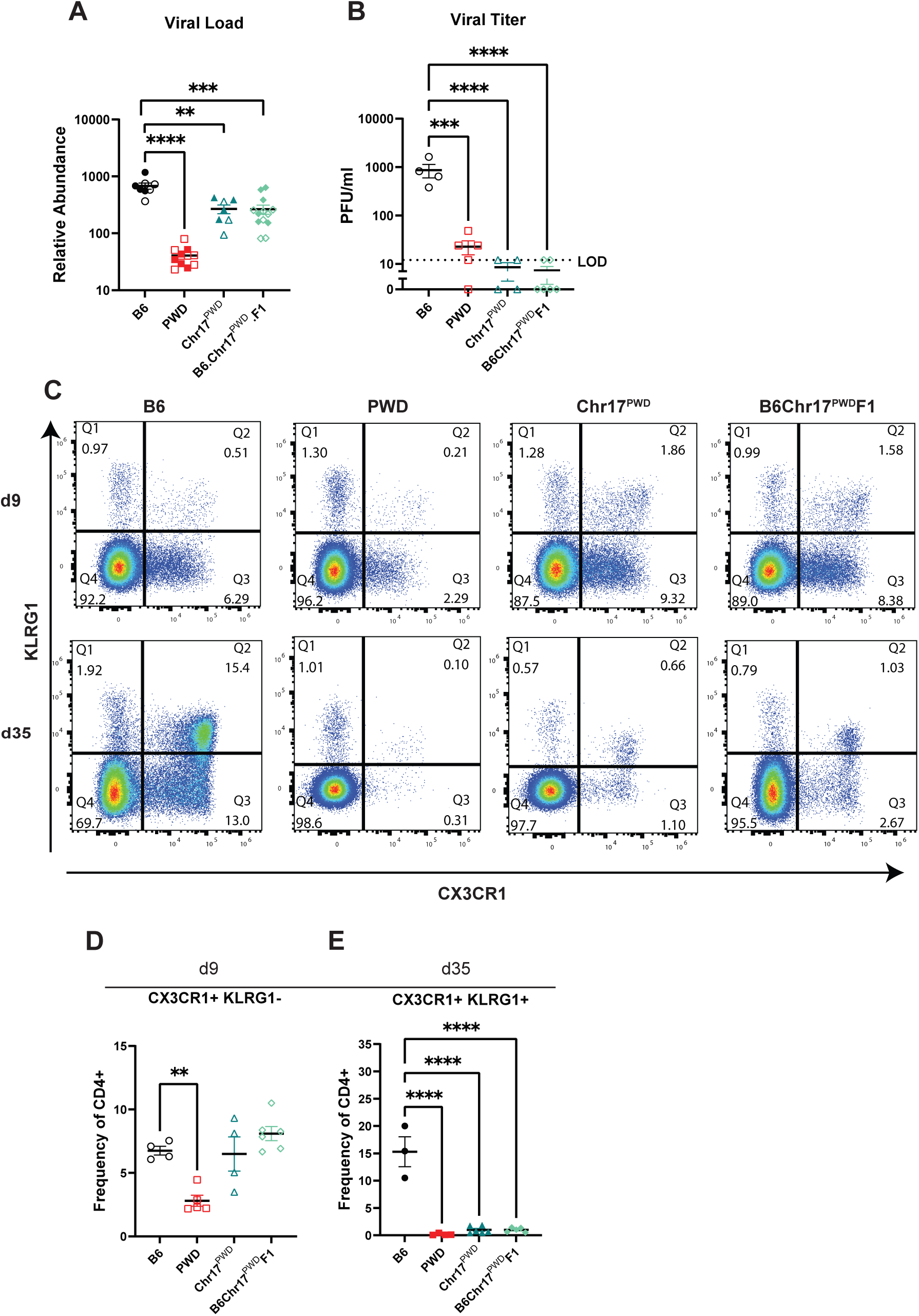
MHV-68 control on PWD chromosome 17 is dominant. Reciprocal F1 crosses between B6 and Chr17PWD was completed. Male and female offspring from the F1 cross, and B6, PWD, and Chr17PWD controls were infected with 105 PFU of MHV-68 via the i.p. route. At d9 and d35 post-infection, spleens were collected followed by assessment of replication of MHV-68 and flow cytometry. Data are shown with sexes pooled. (**A**) Day 9 relative abundance of MHV-68 ORF50 DNA load as assessed by qPCR. (**B**) Comparison of infectious viral titers (expressed as plaque forming units (PFU) per ml) at d9. (**C**) Representative flow cytometric plots of CX3CR1 and KLRG1 expression by CD4+ T cells on d9 and d35 post infection. (**D-E**) Frequency of CX3CR1+ KLRG1- CD4+ T cells and CX3CR1+ KRLG1+ CD4+ T cells (calculated as percent of CD4+ T cells) at d9 and d35 post infection, respectively. For the reciprocal crosses, B6Chr17PWDF1 mice from B6 (female) x Chr17PWD (male) are represented as filled symbols, while Chr17PWDB6F1 mice from B6 (male) x Chr17PWD (female) are represented as open symbols. Only B6Chr17PWDF1 mice were utilized for viral titer in (**B**). PWD and Chr17PWD samples in (**C**) and (**E**) were reproduced from the day 35 time point experiment shown in Fig. 3. Significance of differences between genotypes was determined in (**A** and **E**) by lognormal one-way ANOVA with Dunnett’s multiple comparisons test and in (**D**) by ordinary one-way ANOVA with Dunnett’s multiple comparisons test. Data in (**B**) had a constant of 1 added to them to account for undetectable values. Significance of difference between genotypes was then determined by lognormal one-way ANOVA with Dunnett’s multiple comparisons test. For (**A-B**) and (**D-E**): * indicates p≤0.05, ** indicates p≤0.01, *** indicates p≤0.001, and **** indicates p≤0.0001.

Given the muted cytotoxic T cell response we observed in PWD and Chr17^PWD^ mice (**Fig. 3**), we assessed the frequency of CX3CR1+ KLRG1- and CX3CR1+ KLRG1+ CD4+ T cells in B6Chr17^PWD^F1 mice. We found that B6Chr17^PWD^F1 mice, similar to Chr17^PWD^, have a B6-like expansion of CX3CR1+ KLRG1- CD4+ T cells at d9 (**Fig. 5C-D**). We further observed that B6Chr17^PWD^F1, similar to Chr17^PWD^ and PWD mice, have significantly fewer CX3CR1+ KLRG1+ CD4+ T cells at d35 (**Fig. 5C** and **E**). Taken together, these results demonstrate that B6Chr17^PWD^F1 mice fully phenocopy the T cell response and viral resistance seen in Chr17^PWD^ mice, demonstrating that a single copy of the PWD chromosome 17 is sufficient for both enhanced control of MHV-68 viral burden and muted cytotoxic T cell responses, and suggesting that these two phenotypes are functionally linked.

### Control of viral burden in Chr17^PWD^ mice is NK cell-independent

In our previous study (17), we found that the control of MHV-68 in PWD mice, but not in B6 mice, was NK cell dependent, with an increase in splenic viral load and titer seen after NK depletion. Here, we sought to determine their involvement in control of viral replication in Chr17^PWD^ mice. We initially characterized the frequency of specific subsets of NK cells in B6, PWD, and Chr17^PWD^ mice, using CD11b and CD27 as markers of NK cell maturity (24) at d9, as an acute infection time point where robust differences in viral replication are observed among the strains. We found that the overall frequency of splenic NK cells was comparable between the 3 strains (**Fig. 6A**). However, PWD mice had significantly more CD11b- CD27- (least mature), CD11b- CD27+, and CD11b+ CD27+ NK cell subsets compared to B6 mice, while they also had significantly fewer CD11b+ CD27- (most mature) NK cells (**Fig. 6B-F**). Chr17^PWD^ mice more closely resembled the NK profile of B6 mice, although they had significantly fewer CD11b-CD27+, CD11b+ CD27+ and significantly more CD11b+ CD27- (most mature) NK cells compared to the B6 mice (**Fig. 6B-F**). These results demonstrate that: 1) PWD mice exhibit a unique NK phenotype with higher frequencies of immature subsets, and 2) PWD-encoded genetic variants controlling said phenotype are not captured on chromosome 17.

**Figure 6.**
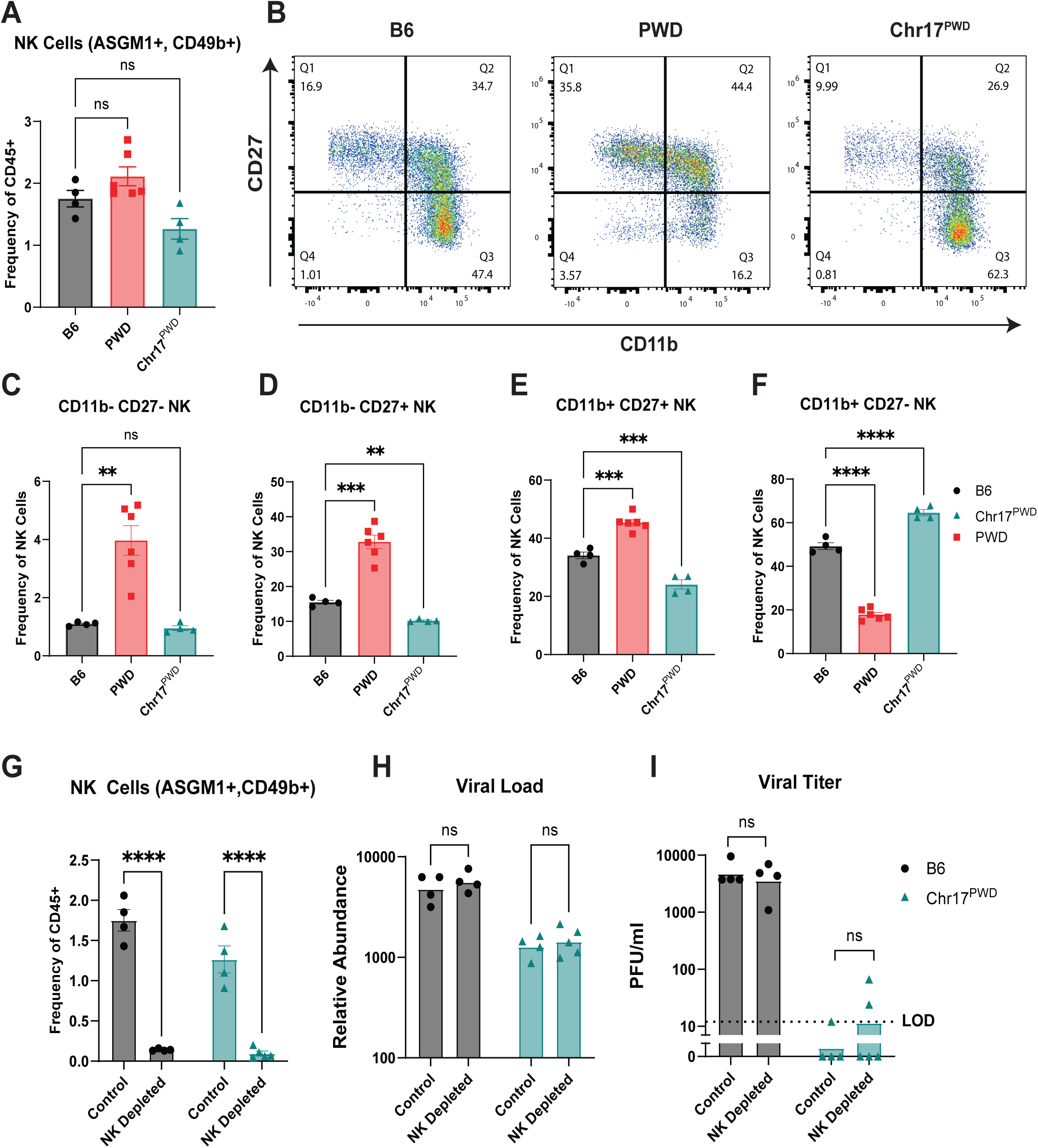
MHV-68 control in Chr17PWD mice is NK cell independent. (**A-F**) Male and female B6, PWD and Chr17PWD mice were infected with 105 pfu of MHV-68 via the i.p. route. Spleens were collected at d9 and flow cytometry was completed to assess immune populations. Male and female data were pooled by genotype. (**A**) Frequency of NK cells (ASGM1+ CD49b+) calculated as percent of CD45+ leukocytes. (**B**) Representative flow plots of CD11b and CD27 expression on NK cells (ASGM1+ CD49b+). (**C-F**) Frequency of NK cell subsets CD11b- CD27-, CD11b- CD27+, CD11b+ CD27+, CD11b+ CD27-, as percent of NK cells (ASGM1+ CD49b+). Significance of differences between genotypes for (**A, E, F**) were determined by one-way ANOVA with Dunnett’s multiple comparisons test. The standard deviations were significantly different, therefore (**C, D**) were analyzed by Brown-Forsythe and Welch ANOVA with Dunnett’s T3 multiple comparisons test. (**G-I**) Male and female B6 and Chr17PWD mice were treated with either anti-NK1.1 or isotype control antibody one day prior to i.p. infection with MHV-68. Spleens were collected at d9 and viral replication and immune phenotypes were assessed. Male and female data were pooled by genotype. (**G**) Frequency of NK cells (ASGM1+ CD49b+) as percent of CD45+ splenocytes. (**H**) Relative abundance of MHV-68 ORF50 DNA load as assessed by qPCR. (**I**) Infectious viral titers (expressed as plaque forming units (PFU) per mL). Significance of differences within strain between NK depleted and isotype controls were determined in (**G**) by two-way ANOVA with an uncorrected Fisher’s LSD, and in (**H**) by two-way ANOVA with Šídák’s multiple comparisons test on the log transformed data. Data in (**I**) had a constant of 1 added and were subsequently log-transformed to account for undetectable values and to ensure that the data is lognormal. Significance of differences within strain between NK depleted and isotype control were then determined by two-way ANOVA with Šídák’s multiple comparisons test on the transformed data. For (**A, C-I**): * indicates p≤0.05, ** indicates p≤0.01, *** indicates p≤0.001, and **** indicates p≤0.0001.

Next, because the muted T cell expansion observed in Chr17^PWD^ mice implied that robust T cell responses are not necessary for control of viral burden, we asked if their resistance to MHV-68 was NK cell-dependent, like that which we reported in PWD mice (17). To this end, we treated B6 and Chr17^PWD^ mice with either anti-NK1.1 or isotype control antibodies one day prior to infection, followed by i.p. MHV-68 infection and assessment of viral replication in the spleen d9 post-infection. We saw a robust depletion of NK cells as detected by flow cytometry in anti-NK1.1-treated animals compared to isotype control, which was comparable in both strains (**Fig. 6G**). However, there was no significant difference in viral load or titer between NK-depleted and isotype control animals for either B6 or Chr17^PWD^ mice (**Fig. 6H** and **I**). Collectively, these results suggest that the PWD-derived loci on chromosome 17 provide resistance to MHV-68 infection in a manner that is NK cell-independent.

### Resistance of PWD mice to MHV-68 is independent of the natural killer cell gene locus on chromosome 6

Compared with PWD, Chr17^PWD^ mice have only an intermediate reduction in viral load, which led us to hypothesize the existence of additional loci that contribute to the PWD MHV-68 resistance. Given our findings that Chr17^PWD^ mice also do not recapitulate the previously identified NK-dependent control of MHV-68 replication (17), we hypothesized the existence of a second MHV-68 resistance locus in the PWD genome mediating these NK-dependent effects. A logical candidate is the highly polymorphic NK cell receptor locus on chromosome 6, which encodes gene variants that profoundly modulate NK cell-mediated killing and recognition of MHC class I alleles by NK cells, especially in the context of betaherpesvirus infection (25). To test this hypothesis, we utilized B6.Chr6^PWD^ (Chr6^PWD^) consomic mice. Because this strain was no longer available in our colony during our initial consomic screen (**Fig. 2**), we cryo-recovered Chr6^PWD^ mice (see Materials and Methods) and, together with B6 and PWD controls, infected them with MHV-68. Spleens were collected at d9, and immune phenotypes were assessed by flow cytometry, and viral replication was assessed by viral load and infectious titer. Surprisingly, Chr6^PWD^ mice exhibited comparable splenic viral loads and titers to B6 mice (**Fig 7A** and **7B**), demonstrating that the NKC^PWD^ locus is not sufficient for enhanced control of viral replication.

**Figure 7.**
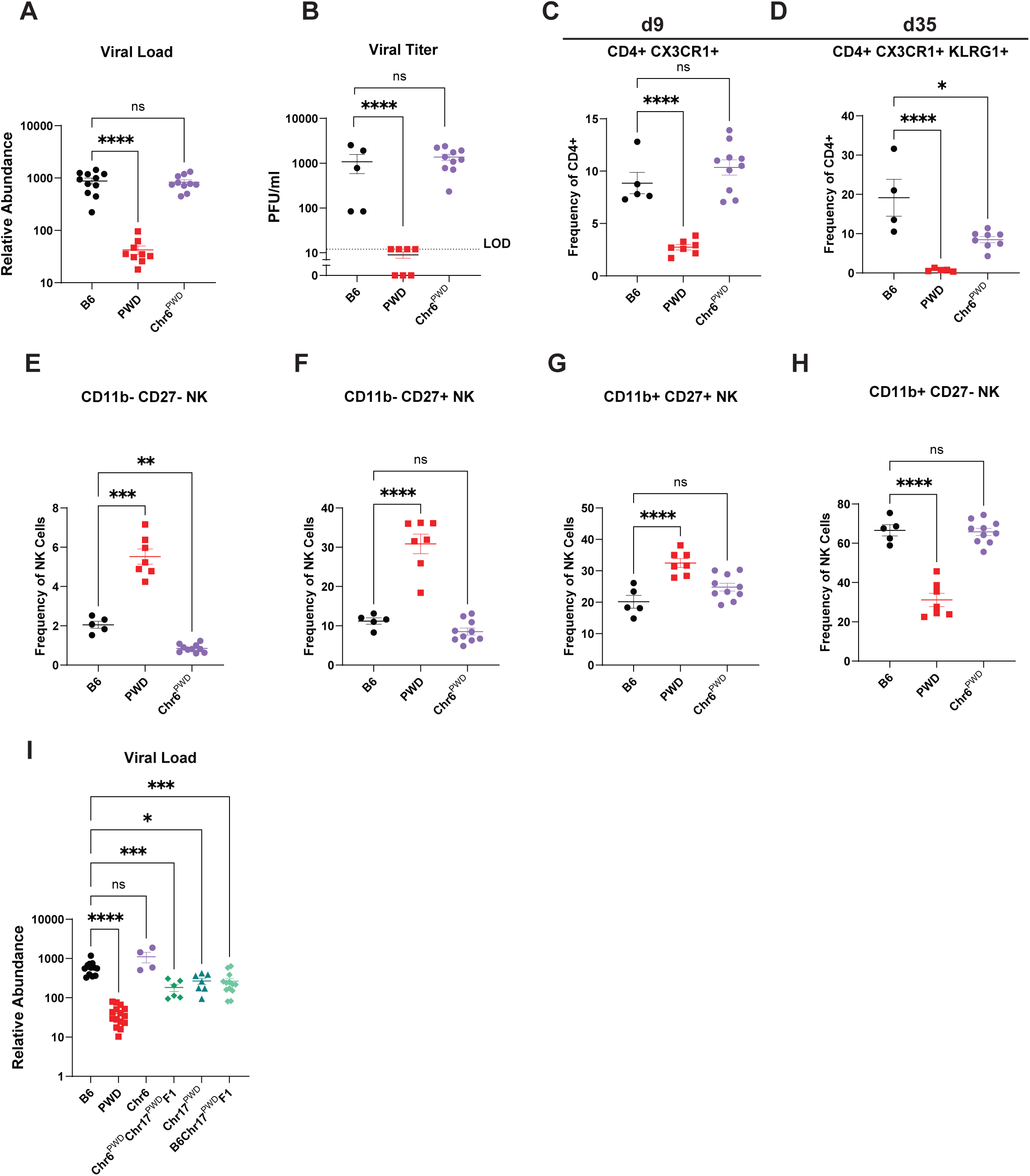
Chr6PWD mice are susceptible to MHV-68. Male and female Chr6PWD mice and B6 and PWD controls were infected with 105 PFU MHV-68 by the i.p. route. Spleens were collected at d9 or d35, and immune phenotypes were assessed by flow cytometry, and viral replication was assessed by viral load and infectious titer. Male and female data were pooled by genotype. (**A**) Relative abundance of MHV-68 ORF50 DNA load at d9 as assessed by qPCR. (**B**) Comparison of infectious viral titers at d9 (expressed as plaque forming units (PFU) per mL). (**C**) and (**D**) Summary graphs of ThCTL, CX3CR1+ KLRG1- CD4+ T cells (d9) and CX3CR1+ KLRG1+ CD4+ T cells (d35) as a frequency of CD4+ cells. (**E-H**) Summary graphs comparing NK cell subsets CD11b- CD27-, CD11b- CD27+, CD11b+ CD27+, CD11b+ CD27- at d9 as frequency of NK cells (ASGM1+ CD49b+). (**I**) Chr6PWDChr17PWDF1 mice were generated by crossing a Chr6PWD (female) with a Chr17PWD (male). Chr6PWDChr17PWDF1 mice and B6 and PWD controls were infected with MHV-68 by the i.p. route. Spleens were collected at d9, relative abundance of MHV-68 ORF50 DNA load was assessed by qPCR. Significance of differences between genotypes was determined in (**A, D,** and **I**) by lognormal one-way ANOVA with Dunnett’s multiple comparisons test except for **I** (Sidak). Data in (**B**) had a constant of 1 added to them to account for undetectable values. Significance of difference between genotypes was then determined by lognormal one-way ANOVA with Dunnett’s multiple comparisons test. Significance of differences in (**C**) was determined by ordinary one-way ANOVA with Dunnett’s multiple comparisons test. Significance of differences between genotypes for (**E-H**) were determined by Brown-Forsythe and Welch ANOVA with Dunnett’s T3 multiple comparisons test. For (**A-I**): * indicates p≤0.05, ** indicates p≤0.01, *** indicates p≤0.001, and **** indicates p≤0.0001.

Our previous data from Chr17^PWD^ and Chr17S^PWD^ mice suggested that the expansion of ThCTL requires a B6 contributed segment at 27.6-49.4 Mb on Chr 17, which correlates with poor viral control. Here, we found that although there was an initial expansion of CX3CR1+ KLRG1- CD4+ T cells at d9 in Chr6^PWD^ mice similar to B6 controls, by d35 the frequency of the CX3CR1+ KLRG1+ CD4+ ThCTL did not reach the level of B6, although it was still much greater compared with PWD mice (**Fig. 7C-D**). These data suggest that although Chr6^PWD^ mice do not control MHV-68 replication better than B6, they do not require the same level of expansion of cytotoxic T cells, possibly due to a differential contribution of NK cells.

We next assessed whether the NKC^PWD^-bearing Chr6^PWD^ mice captured the above-identified immature NK cell phenotype that is present in PWD mice but absent in B6 and Chr17^PWD^ mice (**Fig. 6B-F**). As before, we found that PWD mice had significantly more CD11b-CD27- (least mature), CD11b- CD27+, and CD11b+ CD27+ NK cells compared to B6 mice. Surprisingly we found that Chr6^PWD^ mice, similar to Chr17^PWD^ mice, more closely resembled B6 mice although they had a slight but significant reduction in CD11b- CD27- (least mature) NK cells (**Fig. 7E-H**). Together, these results demonstrate the PWD-derived variants encoded on chromosome 6, which include the NK cell receptor locus, are not sufficient for resistance to MHV-68 and do not regulate NK cell subset frequency.

Given the known allele-specific direct functional interactions of NK cell receptors with MHC(25, 26), we asked if an epistatic interaction between NKC^PWD^ and MHC^PWD^ was required for enhance control of viral replication. To this end, we crossed Chr6^PWD^ and Chr17^PWD^ mice to generate a Chr6^PWD^Chr17^PWD^F1 generation that is heterozygous at both Chr 6 and Chr 17. Male and female F1 mice, along with B6, PWD, and Chr6^PWD^ controls were infected with MHV-68 (i.p.) and spleens were collected at d9 post-infection to assess viral load. We found that Chr6^PWD^Chr17^PWD^F1 mice had an intermediate viral load, comparable to both Chr17^PWD^ and B6Chr17^PWD^F1 mice (**Fig. 7I**), and thus lacking any evidence of additive or synergistic effects between the two chromosomes. These results demonstrate that the NKC^PWD^ locus, at least in a heterozygous state, is not an epistatic modifier of our identified Chr 17 MHV-68 resistance locus.

### Evidence for additional loci in the PWD genome regulating MHV-68 burden

Our initial association mapping findings on viral load in B6PWDF2 mice (**Fig. 1**) suggested the existence of a single partially dominant locus in the PWD genome, which fully controlled viral burden in its homozygous state. In contrast, our physical mapping data on Chr17^PWD^ mice demonstrated the existence of a completely dominant locus, where a single PWD allele was sufficient to provide partial resistance by viral load and full resistance by viral titer (**Fig. 2** and **Fig. 5**). To reconcile these conflicting results, we genotyped the same cohort of B6PWDF2 mice and assessed their viral titers. Segregating this population by their genotype on Chr 17 proximal to our locus of interest (see Methods) yielded two findings. First, mice carrying a homozygous or heterozygous PWD-derived locus of interest on Chr 17 demonstrated PWD-like resistance by viral titer and intermediate resistance by viral load (**Fig. 8A,B**), in agreement with our findings in Chr17^PWD^ mice. However, a large subset of the mice that carried the B6 genotype at the locus of interest on Chr 17 also exhibited PWD-like or intermediate resistance (**Fig. 8A,B**), demonstrating that Chr17^PWD^ is sufficient, but not required for enhanced control of viral burden. To further corroborate this result, we performed a preliminary (given the small sample size) quantitative trait locus (QTL) mapping analysis for viral load and titer in B6PWDF2 mice. This identified several emerging QTL peaks on Chr 6, 15, and 19 (viral load) and Chr 5 and 15 (viral titer), while interestingly no peaks on Chr 17 were observed (**Fig. 8C,D**), likely due to insufficient numbers of mice carrying homozygous (susceptible) B6 alleles on Chr 17 (**Fig. 8A,B**). Collectively, these results suggest the presence of additional non-Chr 17-linked gammaherpesvirus resistance loci in the PWD genome that may regulate control of viral burden by distinct immune mechanisms.

**Figure 8.**
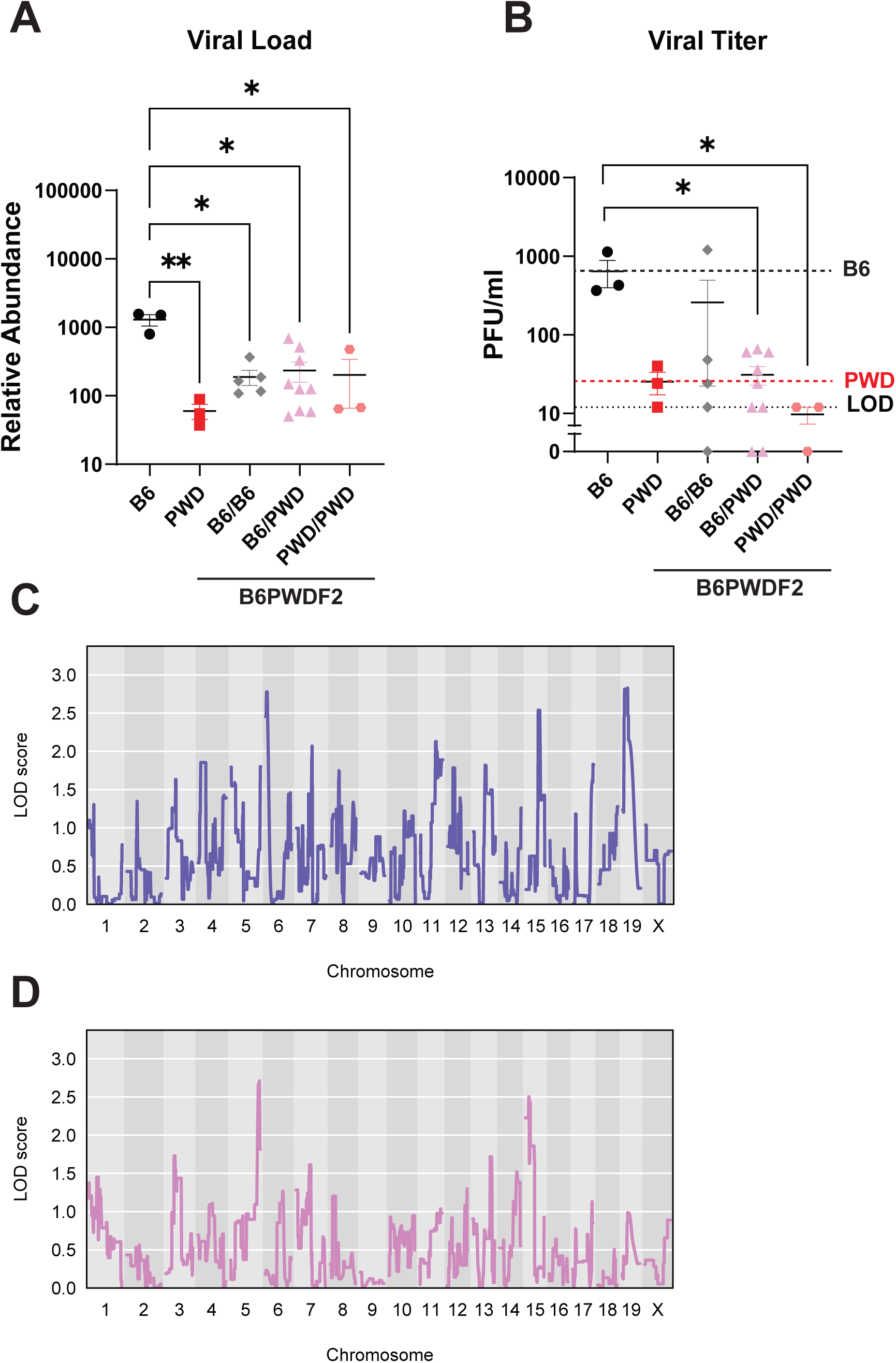
Evidence for additional loci in the PWD genome regulating MHV-68 burden. Male and female B6PWDF2 and control B6 and PWD mice were infected with MHV-68 as in Figure 1. Viral loads as determined by qPCR (**A**) and infectious viral titers (expressed as plaque forming units (PFU) per mL) (**B**). B6PWDF2 mice are stratified by genotype: B6 homozygous (B6/B6), PWD homozygous (PWD/PWD), and heterozygous (B6/PWD) at chromosome 17 34.2-37.8 Mb (H2 region). Significance of differences was determined in (**A**) and (**B**) by lognormal one-way ANOVA with Dunnett’s multiple comparisons test. Data in (**B**) had a constant of 1 added to account for undetectable values, prior to ANOVA. Manhattan plot demonstrating LOD traces for viral load (**C**) and infectious viral titer (**D**). For (**A-B**): * indicates p≤0.05, ** indicates p≤0.01

## Discussion

Previously, we had identified the wild-derived mouse strain PWD as being elite controllers of MHV-68 replication(17). In this study, we sought to determine the host genetic factors contributing to the MHV-68 resistance of PWD mice and investigate the underlying immune mechanisms. We initially found evidence that our resistance phenotype was incompletely dominant, with B6PWDF1 mice demonstrating an intermediate viral load compared with B6 and PWD, and the B6PWDF2 population exhibiting a trimodal distribution. With incomplete dominance, we expect that heterozygous individuals will have an intermediate phenotype compared to the two homozygous parents, which we observed with the B6PWDF1 mice. Similarly, within the B6PWDF2 population, where natural recombination occurs, we expect individuals inheriting the major resistance locus as B6/B6 to be susceptible, those with B6/PWD to show an intermediate phenotype, and PWD/PWD individuals to be resistant, resulting in a trimodal distribution. However, when we performed physical mapping using consomic mice, and identified Chr17^PWD^ mice as having an intermediate viral load, this was inconsistent with an incompletely dominant resistance locus - since under this model homozygous Chr17^PWD^ mice should fully recapitulate the low viral load of PWD mice. Furthermore, our studies of F1 crosses between B6 and Chr17^PWD^ showed that the resistance phenotype linked to Chr 17 is fully dominant, rather than incomplete. These inconsistencies prompted us to perform a genotype-phenotype analysis in the B6PWDF2 population, which demonstrated that a PWD genotype on Chr17 is sufficient, but not required to drive resistance. Taken together, these results suggest that the trimodal phenotype distribution in the B6PWDF2 population arises from multiple genes contributing to the resistance, likely through epistatic interactions, rather than from a single incompletely dominant locus. This sets the stage for future mapping studies to identify additional loci in the PWD genome driving resistance, in particular those that may involve differential contributions of NK cells to the control of viral replication, since this phenotype was not captured in Chr17^PWD^ mice.

In this study, we physically mapped a major gammaherpesvirus resistance locus to PWD chromosome 17, between 27.6-49.4 Mb. Notably, this interval contains the entirety of the MHC/*H-2* region. This finding, together with the findings of EBV GWAS studies (see below), implicates MHC class I and/or class II variants. There are multiple mechanisms by which MHC variants may confer resistance to viral infections, with a major one being differences in antigen presentation that lead to changes in the adaptive response. We found that, consistent with our previously reported findings in PWD mice(17), resistant Chr17^PWD^ mice also lacked a sustained expansion of ThCTL cells and LLEC CD8+ T cells, suggesting that this phenotype may be driven by the MHC. While cytotoxic T cells play an important role in eliminating virally infected cells, there is an important balance between control of viral replication and damage to the host. Infectious mononucleosis is characterized by an overexpansion of CD8+ T cells, leading to splenomegaly, hepatitis, and other sequelae(1). Similarly, in the MHV-68 model of infectious mononucleosis in B6 mice, there is an overexpansion of CD62L- activated CD8+ T cells(27), as well as Vβ4+ CD8+ T cells specific for the viral M1 antigen that are not essential for the control of latent MHV-68(28), and expanded cytotoxic cells can enter tissues such as the liver, where they cause tissue damage resulting in hepatitis(29). Our findings in PWD and Chr17^PWD^ mice thus suggest that sustained expansion of effector cytotoxic T cells may paradoxically indicate inefficient control of gammaherpesvirus. Future studies will need to experimentally test the mechanisms driving resistance of PWD and Chr17^PWD^ mice, including the relative contribution of various T cell subsets (e.g. CD4 and CD8 T cell) or key T cell-derived mediators (e.g. IFNγ), all of which are canonically essential for control of MHV-68 in B6 mice(16). Importantly, while our mapping studies and prior knowledge implicate the MHC locus as a lead candidate, the entirety of our locus of interest on Chr 17 contains 491 protein coding genes, all of which represent potential candidates. Thus, further mapping studies, together with candidate gene and functional studies, are needed to determine the minimal locus required for resistance, identify the causative gene(s) and determine the underlying the mechanism(s).

A second mechanism by which MHC class I variants can impact viral resistance is through direct interactions with natural killer receptors, which are essential for the detection of the “missing-self” in virus-infected cells. NK cells have been shown to be critical for the control of betaherpesviruses HCMV and MCMV, with a marked expansion of NKG2C+ NK cells that generate memory-like responses against these viruses(30, 31). In the context of gammaherpesviruses, patients with NK cell deficiencies fare worse than those with CD8+ T cell deficiencies(32), suggesting NK cells play a critical role in the control of EBV. In previous studies of MHV-68 infection in B6 mice, NK cells were found to be dispensable for control of viral burden(33). However, in our previous study(17), we found that the MHV-68 resistance in PWD mice is at least partially NK cell dependent, recapitulating the essential role of NK cells for control of gammaherpesvirus in humans. Here, we hypothesized that the newly identified resistance locus on Chr 17 could be driven by differential interactions of PWD-derived MHC class I variants with NK cell receptors. Surprisingly, we found that depletion of NK cells had no effect on viral burden in Chr17^PWD^ mice. Given multiple examples of protective MHC-NKC epistatic interactions, including between Ly49P and H-2D^K^ in mice resistant to MCMV(26), we investigated if the NK cell dependence in PWD mice was due to either an NK cell receptor variant on chromosome 6, or due to an epistatic interaction between the PWD-derived NKC and MHC class I loci. Contrary to our expectations, we found no evidence for either hypothesis, as homozygous Chr6^PWD^ mice lacked resistance, and Chr6^PWD^Chr17^PWD^F1 mice did not demonstrate additional resistance compared with Chr17^PWD^ or B6Chr17^PWD^F1 mice. Interestingly, we observed the presence of a unique “immature” CD11b- CD27+ NK cell subset in PWD mice, which was not recapitulated in either Chr6^PWD^ nor Chr17^PWD^ mice. This unique NK cell signature could indicate that the NK cell dependence is instead tied to NK cell function in controlling MHV-68 replication, as other groups have suggested that these subsets may in fact have a “mature” functional capacity with cytotoxic activity and cytokine production(34). Furthermore, a similar subset of early-differentiated NK cells has been hypothesized to be protective against severe EBV infection(35). Further studies could map the loci associated with this NK cell subset, as well as loci driving NK-dependent control of MHV-68 burden.

While Chr17^PWD^ mice fully recapitulate the control of MHV-68 splenic viral titer in PWD mice, they are only partially resistant by DNA viral load. Furthermore, unlike PWD, Chr17^PWD^ mice do not show a reduced frequency of infected (YFP+) splenic B cells compared with B6 mice. This strain is the only one in our study that shows this discordance in viral susceptibility measures. There are two potential explanations for this phenotype: 1) a shift in the kinetics of the virus towards latency earlier in the Chr17^PWD^ mice, or 2) a cellular resistance in a key cell that is preventing productive infection. A host-driven push towards earlier latency would explain the high levels of splenic viral DNA and infected B cells at the peak of lymphoid replication at d9, despite very low viral titers. An interesting caveat to the YFP-bearing MHV-68 is that it cannot differentiate between productive infection (and release of viral particles) vs. latent infection(23). Thus, while we can detect similar frequencies of key B cell subsets that are YFP+ in B6 and Chr17^PWD^ mice, indicating that the B cells in Chr17^PWD^ are not intrinsically resistant to MHV-68 entry (arguing against model #2 above), we are not able to determine if virus is released from these cells. While a mechanistic step-wise assessment of B cell infection is precluded by a lack of *in vitro* B cell infection models for MHV-68, future studies could use *ex vivo* reactivation assays and other measures of MHV-68 latency to further address the possibility of a shift in latency kinetics in Chr17^PWD^ mice, as well as host mechanisms that may drive this phenomenon.

Genetic studies of gammaherpesvirus susceptibility in immunocompetent individuals have been challenging, particularly for primary infection - given the ubiquitous nature of EBV and the fact that the majority of infections are asymptomatic. Initial genome-wide association studies (GWAS) instead relied on surrogate measures, such as serum titer of antibodies to immunodominant EBV antigens like EBNA-1, or electronic health records of infectious mononucleosis. These studies have implicated several MHC class I and II variants as important regulators of these phenotypes, likely through their critical role in shaping T cell-mediated immunity(36–40). Most recently, massive biobank-based GWAS have instead leveraged whole genome sequencing of blood or saliva DNA to quantify reads mapping to the EBV genome, as a surrogate measure of viral load, predominantly during chronic infection. These studies have improved and fine-mapped numerous associations the MHC locus and implicated a number of non-MHC genes(41–43). While these studies are exciting, they are far from definitive. First, they predominantly assess the chronic phase of viral replication, while initial control of acute replication is what likely sets the stage for subsequent immune (dys)regulation(44). Second, these studies typically detected EBV “DNAemia” at very low levels (1-3 reads per individual) and only in about 10% of individuals, and thus they miss the viral dynamics in the majority of the ∼90% of infected individuals. In this light, our own findings in a mouse model of gammaherpesvirus infection, which allows for temporal control of infection, causative linkage assessment, and mechanistic experimentation, are fully congruent with (and complementary to) the human studies, strongly implicating the MHC locus and additional variants elsewhere in the genome. Our findings also strongly implicate T cell immunity as a major MHC-dependent driver of susceptibility, but with a surprisingly inverse relationship between the magnitude of expansion of effector T cell responses and viral control. Future studies will allow for further mechanistic assessment of this link, and deeper functional comparisons to the emerging powerful human GWAS.

## Material and Methods

### Animals

B6, PWD, and B6.Chr^PWD^ consomic mice were purchased from The Jackson Laboratory (Bar Harbor, ME), then bred and housed in a single room within the vivarium at the University of Vermont for ten or more generations prior to experimentation. To ensure their correct identity, and to enhance rigor and reproducibility of these studies, B6.Chr^PWD^ consomic mice were subjected to genome-wide SNP genotyping using DartMouse genotyping services (Dartmouth College, NH USA). All mice used in this study were of the expected genotypes, with the following exception. Chr17S^PWD^ mice were found to carry a homozygous B6-derived interval between 30 – 45 Mb on Chr 17, encompassing *H2*, as we have previously reported (19).

Chr17^PWD^ mice were provided by Dr. Jiri Forejt (Institute of Molecular Genetics of the Czech Academy of Sciences, Prague, Czech Republic). To ensure their correct identity, these mice were subjected to genome-wide SNP genotyping using Transnetyx mini mouse universal genotyping array (MiniMUGA) and had the expected genotype. B6, PWD, and Chr17S^PWD^ mice were concurrently subjected to genome-wide SNP genotyping using the Transnetyx MiniMUGA, allowing us to refine the Chr17S^PWD^ homozygous B6-derived interval to 27.6-49.4 Mb on Chr 17. This B6-derived interval contains the entire MHC/*H2* at ∼34.2-37.8 Mb.

Chr6^PWD^ mice were cryorecovered by Dr. Jason Bubier (Jackson Laboratories, Bar Harbor, Maine) and shipped to the University of Vermont. To ensure their correct identity, and to enhance rigor and reproducibility of these studies, Chr6^PWD^ mice were subjected to genome-wide SNP genotyping using Transnetyx MiniMUGA and had the expected genome. These mice were then bred and housed in the same room as the other mice in this study. These mice were bred for one generation in our vivarium prior to experimentation.

F1 and F2 intercrosses between strains were generated as follows. A PWD male was mated with a B6 female and the resulting B6PWDF1 progeny were intercrossed to produce B6PWDF2 intercross progeny. B6 and Chr17^PWD^ mice were mated in reciprocal crosses to generate B6Chr17^PWD^F1 and Chr17^PWD^B6F1 progeny. Chr6^PWD^ females were mated with a Chr17^PWD^ male to generate Chr6^PWD^Chr17^PWD^F1 progeny. The experimental procedures used in this study were approved by the Animal Care and Use Committee of the University of Vermont.

### Viruses

MHV-68 (clone G2.4) was provided by Dr. Edward J. Usherwood (The Geisel School of Medicine at Dartmouth College, Hanover, NH). The virus was propagated by infection of 3T3 cells at a multiplicity of infection (MOI) of 0.1, as previously described (45, 46). Infectious virus was measured via plaque titration by infecting confluent 3T3 cells using 1:10 serial dilutions of viral inoculate in DMEM (Cytiva Life Sciences) plus 10% (v/v) FBS (Fisher Scientific) and 5% (v/v) penicillin/streptomycin (P/S: Cytiva Life Sciences) at 37°C for 1 h, after which the cells were overlaid with 1.95% (w/v) solution of carboxymethylcellulose (Sigma-Aldrich) in DMEM plus 10% (v/v) FBS and 5% (v/v) penicillin/streptomycin and incubated for 7 days. After incubation, cells were fixed with methanol and stained with 0.05% crystal violet (w/v) (Fisher Scientific) in 20% (v/v) methanol, and then the plaques were enumerated to determine titer (PFU/ml).

MHV68-histone 2B-YFP (23) was provided by Dr. Edward J. Usherwood with permission by Dr. Samuel Speck (Professor Emeritus, Emory Vaccine Center, Atlanta, GA). The virus was propagated as described above for the WT MHV-68 virus.

### MHV-68 infections

B6, PWD, and B6.Chr^PWD^ mice for the consomic screen were infected with MHV-68 at an average of 33 weeks of age (range, 6-51 wk). For subsequent experiments, B6, PWD, B6.Chr^PWD^, B6PWDF1, B6PWDF2, B6Chr17^PWD^F1, and Chr6^PWD^Chr17^PWD^F1 mice were infected with MHV-68 at an average of 11 weeks of age (range, 8-30 wk). All MHV-68 infections were administered via intraperitoneal injection of 10^5^ PFU of either WT MHV-68 or MHV68-H2bYFP in a total volume of 100 ul of PBS to allow for viral replication and latency in the spleen that is fully established by ∼1 month post-infection (47). Mice were euthanized at 9, 16, or 35 days post-infection, and spleen was collected and processed as described below for each downstream assay.

### Measurement of infectious virus titer

Infectious viral titers in the spleen were determined by plaque assay following a modified procedure to the one described above for initial infectious viral titer and as previously described (17). Briefly, spleens were frozen at −80°C in 600 ul of media used for plaque assays (DMEM+10% [v/v] FBS+P/S) prior to the assay. Samples were thawed and virus was released via sonication and clarified via centrifugation. 1:10 serial dilutions of the resulting sample viral suspensions were added in 0.5-ml volumes to 3T3 cell monolayers and incubated at 37°C for 1 h to allow the virus to absorb. After the incubation, viral suspensions were removed from the monolayers before overlaying with 1.95% (w/v) solution of carboxymethylcellulose in DMEM plus 10% (v/v) FBS + P/S. After 7 d of incubation at 37°C cells were fixed with methanol and stained with crystal violet, and then the plaques were enumerated to calculate PFU/ml.

### Measurement of viral DNA load

Viral DNA was quantified via quantitative PCR (qPCR) using a modified approach from what was previously described (17). DNA was extracted from spleen tissue using a Qiagen DNeasy blood and tissue kit. 10 mg of spleen tissue were homogenized with a metal bead and bead beater for 1 minute in lysis buffer, prior to proceeding with the manufacturer’s instructions. Concentration was determined via Nanodrop (Thermo Scientific Nanodrop 2000 spectrophotomer) and samples were normalized to 35 ng/ul prior to qPCR.

qPCR was performed according to the manufacturer’s instructions using the PowerTrack SYBR Green Kit (Applied Biosystems) and 100 nM primers complementary to the sequence of the MHV-68 ORF50 gene (48, 49) and samples were run on the QuantStudio 3 real-time PCR system (Applied Biosystems). MHV-68 ORF50 gene abundance was normalized by the abundance of the cellular DNA reference gene *Mapk14* and calculated by a comparative cycle threshold (Ct) method (2^−ΔCt^) and multiplied by a factor of 10,000 for ease of visualization, that is, relative abundance = 2^−(MHV-68 ORF50 Ct − Mapk14 Ct)^ × 10,000. Relative abundance was not further normalized against a reference control experimental group.

### Flow Cytometry

At 9, 16, and 35 days post-infection, mice were euthanized and spleens were collected and processed for staining for flow cytometry, as previously described (17). Briefly, spleens were mechanically dissociated by a syringe plunger between two pieces of nylon 200-micron mesh, until a single cell suspension was created. Resulting cell suspensions underwent RBC lysis by incubation in 0.8% ammonium chloride solution (STEMCELL Technologies).

For flow cytometry analysis of surface staining with permeabilization, cells were stained with UV-Blue LIVE/DEAD fixable stain (Invitrogen) and then surface stained for the following markers: CD45, CD19 (Invitrogen), CD11b, TCRβ, CD4, CD8a, CX3CR1, anti-asialo GM1 (ASGM1), NK1.1 (Biolegend), KLRG1, CD49b, and CD27 (BD Biosciences). Cells were then fixed, permeabilized, and stained for GZMB (BioLegend). Surface staining was completed in brilliant stain buffer (BD Biosciences).

For flow cytometry staining to assess B cells, cells were stained with UV-Blue LIVE/DEAD fixable stain (Invitrogen) and surface stained for the following markers: CD45, CD23, CD19 (Invitrogen), TCRβ, CD95, IgM, CD80, IgD, CD43, CD93, CD45R, GL7, and CD138 (BioLegend). Cells were then fixed 1% paraformaldehyde (Sigma-Aldrich).

All stained cells were analyzed on a Cytek Aurora (Cytek Biosciences). Flow cytometry data analysis was performed using FlowJo software versions 10.10 (BD Biosciences).

### In vivo NK depletion

On the day prior to MHV-68 infection (day −1), B6 and Chr17^PWD^ mice were i.p. injected with 0.1 mg of anti-NK1.1 (clone PK136; BioLegend) or mouse IgG2a, κ isotype control (BioLegend), delivered in a total volume of 100 µl of PBS. NK cell depletion was verified by flow cytometry analysis of CD19^-^TCRβ^−^CD4^−^CD8^−^CD49b^+^ NK1.1^+^ cell populations.

### QTL Mapping

B6PWDF2 mice were genotyped by miniMUGA, as described above. Raw SNP calls were assigned parent genotypes using the R package, stuart(50), prior to QTL mapping. To map genetic variants associated with viral load and viral titer, the R package, R/qtl2(51) was utilized. Viral load and viral titer were determined as described above. For both measures, data from individual mice were utilized, with age and sex as covariates. For viral load, a log transformation was performed, and for viral titer 1 was added to all values to remove zero values before log transformation. Thresholds for 10% and 20% genome-wide significance were generated utilizing 1,000 permutations.

### Statistical Analysis

Statistical analyses were carried out using GraphPad Prism software versions 10.0.2-10.6. Details of the analyses are provided in the figure legends, including the specific tests used to assess the significance of the observed differences and indication of adjustments for multiple comparisons when appropriate. Comparisons were assessed for effects between genotypes (B6, PWD, B6.Chr^PWD^, and genetic crosses) as indicated. All center values represent the mean, and error bars represent the SEM. A *p* value <0.05 was considered significant. Comparisons are indicated by brackets, and *p* values are noted as follows: \**p* ≤ 0.05, \*\**p* ≤ 0.01, \*\*\**p* ≤ 0.001, and \*\*\*\**p* ≤ 0.0001. A lack of an indicated *p* value signifies a lack of significant difference.

## Acknowledgements

The authors would like to acknowledge Dr. Samuel Speck for the generation and sharing of the MHV68-histone 2B-YFP virus and Dr. Jiri Forjet for providing Chr17^PWD^ mice to establish a colony. The authors would also like to acknowledge the Flow Cytometry and Cell Sorting facility at the University of Vermont Larner College of Medicine for the use of their facilities and resources.

---

**Supplemental 1.**
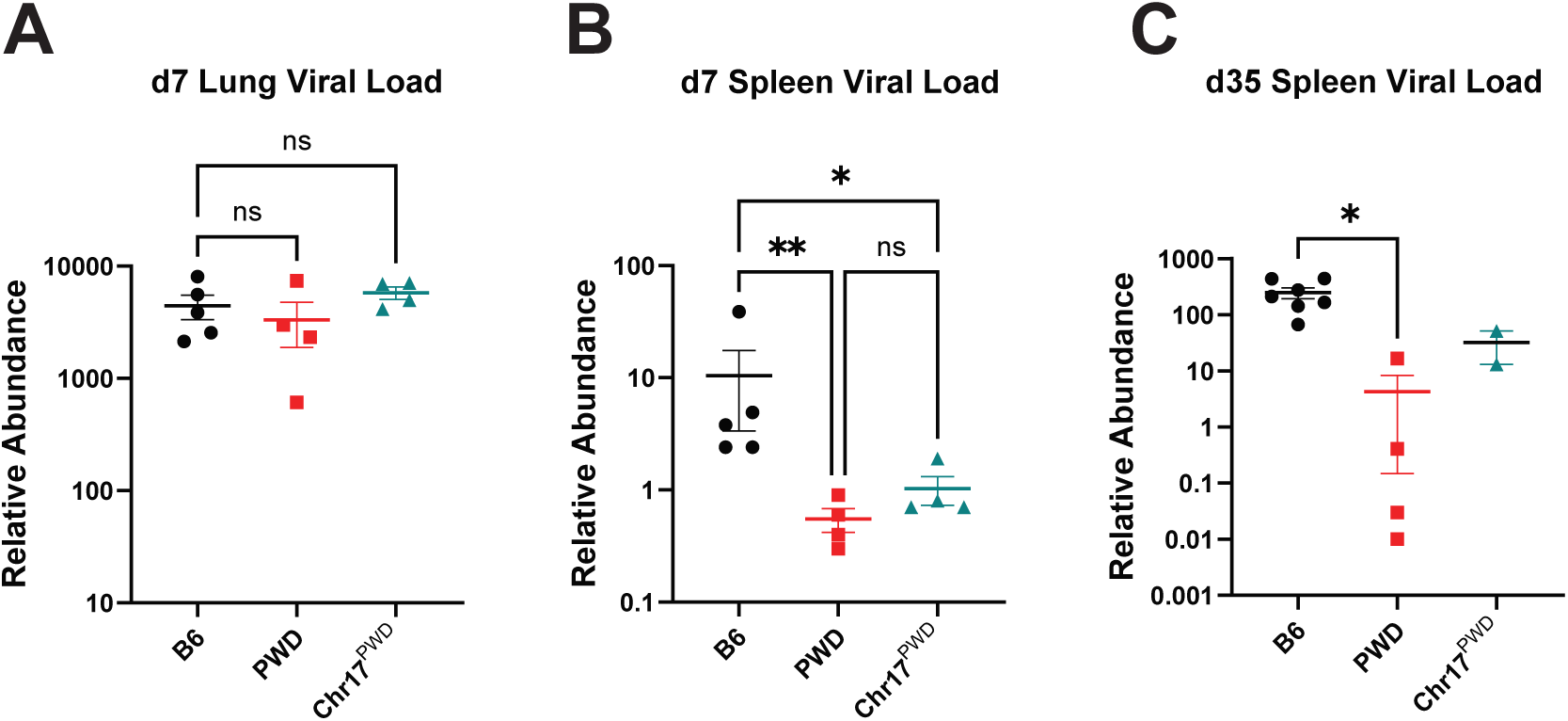
Intranasal MHV-68 infection. Male and female Chr17PWD, B6, and PWD mice were infected with 104 PFU of MHV-68 via the intranasal route. At days 9 and 35 post-infection, lungs (d9 only) and spleens were collected and replication of MHV-68 was assessed. Male and female data were pooled by genotype. (**A-B**) Relative abundance of MHV-68 DNA load, as assessed by qPCR for the ORF50 gene in lungs and spleens at day 7, respectively. (**C**) Relative abundance of MHV-68 DNA load, as assessed by qPCR for the ORF50 gene in spleens at day 35. Significance of differences between genotypes in (**A-C**) were determined by lognormal one-way ANOVA with Tukey’s multiple comparisons test, except in **A** (Dunnett’s) and **C** (Brown-Forsythe and Welch). For (**A-C**): * indicates p≤0.05 ** indicates p≤0.01.

**Supplemental 2:**
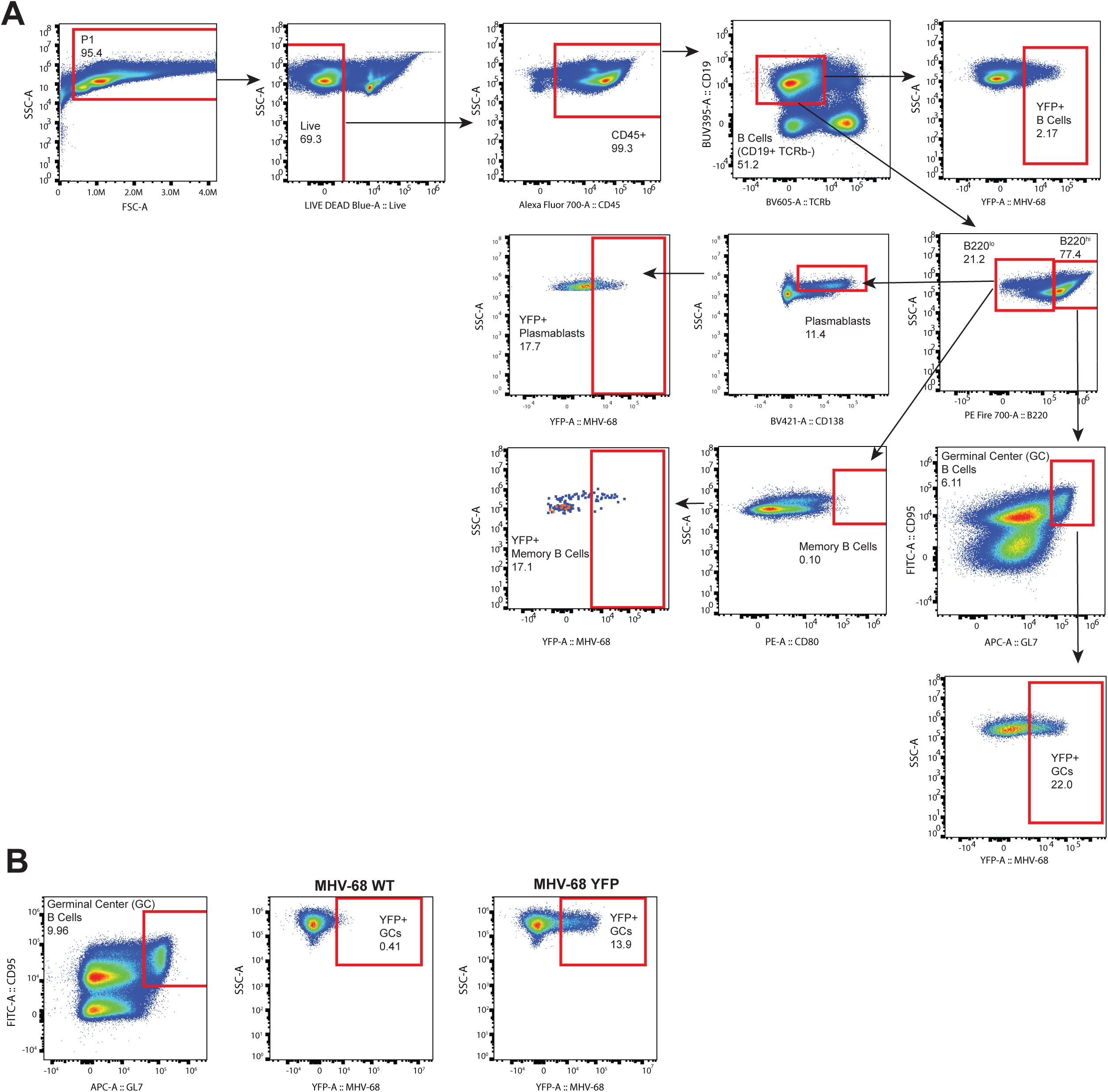
Flow cytometry gating strategy for B cell subsets. Mice were infected with 105 PFU of a YFP reporter-expressing MHV-68 or MHV-68 WT (B only) by the i.p. route. Spleens were collected at d9 post infection and flow cytometry was completed to assess B cell subset frequencies and YFP+ frequency. (**A**) The gating strategy for analysis was completed with the red boxes representing populations of interest. (**B**) Gating strategy for germinal center B cells (GL7+ CD95+) and YFP+ GCs based on MHV-68 WT infected mice.

## References

1. Damania B, Kenney SC, Raab-Traub N. 2022. Epstein-Barr virus: Biology and clinical disease. Cell 185:3652–3670.

2. Jouanguy E, Béziat V, Mogensen TH, Casanova J-L, Tangye SG, Zhang S-Y. 2020. Human inborn errors of immunity to herpes viruses. Current Opinion in Immunology 62:106–122.

3. Hjalgrim H, Askling J, Rostgaard K, Hamilton-Dutoit S, Frisch M, Zhang J-S, Madsen M, Rosdahl N, Konradsen HB, Storm HH, Melbye M. 2003. Characteristics of Hodgkin’s Lymphoma after Infectious Mononucleosis. New England Journal of Medicine 349:1324–1332.

4. Ascherio A, Munger KL. 2007. Environmental risk factors for multiple sclerosis. Part I: The role of infection. Annals of Neurology 61:288–299.

5. Bjornevik K, Cortese M, Healy BC, Kuhle J, Mina MJ, Leng Y, Elledge SJ, Niebuhr DW, Scher AI, Munger KL, Ascherio A. 2022. Longitudinal analysis reveals high prevalence of Epstein-Barr virus associated with multiple sclerosis. Science 375:296–301.

6. Houen G, Trier NH. 2021. Epstein-Barr Virus and Systemic Autoimmune Diseases. Frontiers in Immunology 11.

7. Jog NR, Young KA, Munroe ME, Harmon MT, Guthridge JM, Kelly JA, Kamen DL, Gilkeson GS, Weisman MH, Karp DR, Gaffney PM, Harley JB, Wallace DJ, Norris JM, James JA. 2019. Association of Epstein-Barr virus serological reactivation with transitioning to systemic lupus erythematosus in at-risk individuals. Annals of the Rheumatic Diseases 78:1235–1241.

8. Jog NR, James JA. 2021. Epstein Barr Virus and Autoimmune Responses in Systemic Lupus Erythematosus. Frontiers in Immunology 11.

9. Younis S, Moutusy SI, Rasouli S, Jahanbani S, Pandit M, Wu X, Acharya S, Sharpe O, Wijeratne TU, Harris ML, Yang EY, Chaichian Y, Parsafar S, Baker MC, Harley JB, Meffre E, Steinman L, Marshak-Rothstein A, James JA, Martinez OM, Utz PJ, Orange DE, Lanz TV, Robinson WH. 2025. Epstein-Barr virus reprograms autoreactive B cells as antigen-presenting cells in systemic lupus erythematosus. Science Translational Medicine 17:eady0210.

10. Yu H, Robertson ES. 2023. Epstein-Barr Virus History and Pathogenesis. Viruses 15.

11. Münz C. 2017. Epstein–Barr Virus-Specific Immune Control by Innate Lymphocytes. Frontiers in Immunology 8.

12. Orange JS. 2013. Natural killer cell deficiency. Journal of Allergy and Clinical Immunology 132:515–525.

13. Cesarman E. 2014. Gammaherpesviruses and Lymphoproliferative Disorders. Annual Review of Pathology: Mechanisms of Disease 9:349–372.

14. Agostini S, Mancuso R, Guerini FR, D’Alfonso S, Agliardi C, Hernis A, Zanzottera M, Barizzone N, Leone MA, Caputo D, Rovaris M, Clerici M. 2018. HLA alleles modulate EBV viral load in multiple sclerosis. Journal of Translational Medicine 16:80.

15. Hedström AK, Huang J, Michel A, Butt J, Brenner N, Hillert J, Waterboer T, Kockum I, Olsson T, Alfredsson L. 2020. High Levels of Epstein–Barr Virus Nuclear Antigen-1-Specific Antibodies and Infectious Mononucleosis Act Both Independently and Synergistically to Increase Multiple Sclerosis Risk. Frontiers in Neurology 10.

16. Wang Y, Tibbetts SA, Krug LT. 2021. Conquering the Host: Determinants of Pathogenesis Learned from Murine Gammaherpesvirus 68. Annual Review of Virology 8:349–371.

17. Holt EA, Waytashek CM, Sessions KJ, Asarian L, Lahue KG, Usherwood EJ, Teuscher C, Krementsov DN. 2023. Host Genetic Variation Has a Profound Impact on Immune Responses Mediating Control of Viral Load in Chronic Gammaherpesvirus Infection. The Journal of Immunology 211:1526–1539.

18. Gregorová S, Divina P, Storchova R, Trachtulec Z, Fotopulosova V, Svenson KL, Donahue LR, Paigen B, Forejt J. 2008. Mouse consomic strains: exploiting genetic divergence between Mus m. musculus and Mus m. domesticus subspecies. Genome Res 18:509–15.

19. Lahue KG, Lara MK, Linton AA, Lavoie B, Fang Q, McGill MM, Crothers JW, Teuscher C, Mawe GM, Tyler AL, Mahoney JM, Krementsov DN. 2020. Identification of novel loci controlling inflammatory bowel disease susceptibility utilizing the genetic diversity of wild-derived mice. Genes Immun 21:311–325.

20. Renkema KR, Huggins MA, Borges da Silva H, Knutson TP, Henzler CM, Hamilton SE. 2020. KLRG1(+) Memory CD8 T Cells Combine Properties of Short-Lived Effectors and Long-Lived Memory. J Immunol 205:1059–1069.

21. Callan MFC, Steven N, Krausa P, Wilson JDK, Moss PAH, Gillespie GM, Bell JI, Rickinson AB, McMichael AJ. 1996. Large clonal expansions of CD8+ T cells in acute infectious mononucleosis. Nature Medicine 2:906–911.

22. Stevenson PG, Belz GT, Altman JD, Doherty PC. 1999. Changing patterns of dominance in the CD8+ T cell response during acute and persistent murine γ-herpesvirus infection. European Journal of Immunology 29:1059–1067.

23. Collins CM, Speck SH. 2012. Tracking Murine Gammaherpesvirus 68 Infection of Germinal Center B Cells In Vivo. PLOS ONE 7:e33230.

24. Chiossone L, Chaix J, Fuseri N, Roth C, Vivier E, Walzer T. 2009. Maturation of mouse NK cells is a 4-stage developmental program. Blood 113:5488–5496.

25. Hisashi Arase ESM, Ann E. Campbell, Ann B. Hill, Lewis L. Lanier. 2002. Direct Recognition of Cytomegalovirus by Activating and Inhibitory NK Cell Receptors. Science 296:1323–1326.

26. Martin MP, Gao X, Lee J-H, Nelson GW, Detels R, Goedert JJ, Buchbinder S, Hoots K, Vlahov D, Trowsdale J, Wilson M, O’Brien SJ, Carrington M. 2002. Epistatic interaction between KIR3DS1 and HLA-B delays the progression to AIDS. Nature Genetics 31:429–434.

27. Tripp RA, Hamilton-Easton AM, Cardin RD, Nguyen P, Behm FG, Woodland DL, Doherty PC, Blackman MA. 1997. Pathogenesis of an infectious mononucleosis-like disease induced by a murine gamma-herpesvirus: role for a viral superantigen? J Exp Med 185:1641–50.

28. Flaño E, Hardy CL, Kim IJ, Frankling C, Coppola MA, Nguyen P, Woodland DL, Blackman MA. 2004. T cell reactivity during infectious mononucleosis and persistent gammaherpesvirus infection in mice. J Immunol 172:3078–85.

29. Kanai K, Kageyama S, Yoshie O. 2023. Involvement of TLR4 in Acute Hepatitis Associated with Airway Infection of Murine γ-Herpesvirus 68. The Journal of Immunology 211:1550–1560.

30. Sun JC, Beilke JN, Lanier LL. 2009. Adaptive immune features of natural killer cells. Nature 457:557–561.

31. Gumá Mn, Angulo A, Vilches C, Gómez-Lozano N, Malats Nr, López-Botet M. 2004. Imprint of human cytomegalovirus infection on the NK cell receptor repertoire. Blood 104:3664–3671.

32. Chijioke O, Landtwing V, Münz C. 2016. NK Cell Influence on the Outcome of Primary Epstein–Barr Virus Infection. Frontiers in Immunology 7.

33. Usherwood EJ, Meadows SK, Crist SG, Bellfy SC, Sentman CL. 2005. Control of murine gammaherpesvirus infection is independent of NK cells. European Journal of Immunology 35:2956–2961.

34. Hayakawa Y, Huntington ND, Nutt SL, Smyth MJ. 2006. Functional subsets of mouse natural killer cells. Immunol Rev 214:47–55.

35. Azzi T, Lünemann A, Murer A, Ueda S, Béziat V, Malmberg K-J, Staubli G, Gysin C, Berger C, Münz C, Chijioke O, Nadal D. 2014. Role for early-differentiated natural killer cells in infectious mononucleosis. Blood 124:2533–2543.

36. Sallah N, Miley W, Labo N, Carstensen T, Fatumo S, Gurdasani D, Pollard MO, Dilthey AT, Mentzer AJ, Marshall V, Cornejo Castro EM, Pomilla C, Young EH, Asiki G, Hibberd ML, Sandhu M, Kellam P, Newton R, Whitby D, Barroso I. 2020. Distinct genetic architectures and environmental factors associate with host response to the gamma2-herpesvirus infections. Nat Commun 11:3849.

37. Sallah N, Carstensen T, Wakeham K, Bagni R, Labo N, Pollard MO, Gurdasani D, Ekoru K, Pomilla C, Young EH, Fatumo S, Asiki G, Kamali A, Sandhu M, Kellam P, Whitby D, Barroso I, Newton R. 2017. Whole-genome association study of antibody response to Epstein-Barr virus in an African population: a pilot. Glob Health Epidemiol Genom 2:e18.

38. Rubicz R, Yolken R, Drigalenko E, Carless MA, Dyer TD, Bauman L, Melton PE, Kent JW, Jr., Harley JB, Curran JE, Johnson MP, Cole SA, Almasy L, Moses EK, Dhurandhar NV, Kraig E, Blangero J, Leach CT, Göring HH. 2013. A genome-wide integrative genomic study localizes genetic factors influencing antibodies against Epstein-Barr virus nuclear antigen 1 (EBNA-1). PLoS Genet 9:e1003147.

39. Jiang L, Kerchberger VE, Shaffer C, Dickson AL, Ormseth MJ, Daniel LL, Leon BGC, Cox NJ, Chung CP, Wei WQ, Stein CM, Feng Q. 2022. Genome-wide association analyses of common infections in a large practice-based biobank. BMC Genomics 23:672.

40. Tian C, Hromatka BS, Kiefer AK, Eriksson N, Noble SM, Tung JY, Hinds DA. 2017. Genome-wide association and HLA region fine-mapping studies identify susceptibility loci for multiple common infections. Nat Commun 8:599.

41. Schmidt A, Alawathurage TM, David FS, Ogawa Y, Frach L, Richter S, Schaefer M, Mathey CM, Henne SK, Nagao G, Tanaka H, Azekawa S, Lee K, Fukunaga N, Hamamoto J, Kabata H, Masaki K, Kamata H, Ikemura S, Chubachi S, Okamori S, Terai H, Morita A, Asakura T, Ishii M, Fukunaga K, Uwamino Y, Uchida S, Uno S, Nishimura T, Hasegawa N, Yanagita E, Nishihara H, Sasaki J, Morisaki H, Sato T, Kitagawa Y, Matsubara Y, Mikami Y, Nanki K, Kanai T, Edahiro R, Shirai Y, Sonehara K, Okuzaki D, Motooka D, Kanai M, Naito T, Yamamoto K, Wang QS, et al. 2026. Host control of persistent Epstein–Barr virus infection. Nature doi:10.1038/s41586-026-10274-4.

42. Nyeo SS, Cumming EM, Burren OS, Pagadala MS, Gutierrez JC, Ali TA, Kida LC, Chen Y, Chu H, Hu F, Zou XZ, Hollis B, Fabre MA, MacArthur S, Wang Q, Ludwig LS, Dey KK, Petrovski S, Dhindsa RS, Lareau CA. 2026. Population-scale sequencing resolves determinants of persistent EBV DNA. Nature 650:664–672.

43. Kamitaki N, Tang D, McCarroll SA, Loh PR. 2026. The DNA virome varies with human genes and environments. Nature doi:10.1038/s41586-026-10288-y.

44. Murdock SJ, Owens SM, Oldenburg DG, Forrest JC. 2026. The amplitude of gammaherpesvirus lytic replication dictates adaptive immune activation: Potential implications for KSHV LANA in immune evasion. bioRxiv doi:10.64898/2026.02.05.703987.

45. Obar JJ, Crist SG, Gondek DC, Usherwood EJ. 2004. Different functional capacities of latent and lytic antigen-specific CD8 T cells in murine gammaherpesvirus infection. J Immunol 172:1213–9.

46. Sunil-Chandra NP, Efstathiou S, Arno J, Nash AA. 1992. Virological and pathological features of mice infected with murine gamma-herpesvirus 68. J Gen Virol 73 (Pt 9):2347–56.

47. Casiraghi C, Shanina I, Cho S, Freeman ML, Blackman MA, Horwitz MS. 2012. Gammaherpesvirus Latency Accentuates EAE Pathogenesis: Relevance to Epstein-Barr Virus and Multiple Sclerosis. PLOS Pathogens 8:e1002715.

48. Usherwood EJ, Ward KA, Blackman MA, Stewart JP, Woodland DL. 2001. Latent antigen vaccination in a model gammaherpesvirus infection. J Virol 75:8283–8.

49. Virgin HW, Latreille P, Wamsley P, Hallsworth K, Weck KE, Canto AJD, Speck SH. 1997. Complete sequence and genomic analysis of murine gammaherpesvirus 68. Journal of Virology 71:5894–5904.

50. Bourdon M, Montagutelli X. 2022. stuart: an R package for the curation of SNP genotypes from experimental crosses. G3 (Bethesda) 12.

51. Broman KW, Gatti DM, Simecek P, Furlotte NA, Prins P, Sen Ś, Yandell BS, Churchill GA. 2019. R/qtl2: Software for Mapping Quantitative Trait Loci with High-Dimensional Data and Multiparent Populations. Genetics 211:495–502.

